# A *pals-25* gain-of-function allele triggers systemic resistance against natural pathogens of *C. elegans*

**DOI:** 10.1101/2022.06.28.497889

**Authors:** Spencer S. Gang, Manish Grover, Kirthi C. Reddy, Deevya Raman, Ya-Ting Chang, Damian C. Ekiert, Michalis Barkoulas, Emily R. Troemel

## Abstract

Regulation of immunity throughout an organism is critical for host defense. Previous studies in the nematode *Caenorhabditis elegans* have described an “ON/OFF” immune switch comprised of the antagonistic paralogs PALS-25 and PALS-22, which regulate resistance against intestinal and epidermal pathogens. Here, we identify and characterize a PALS-25 gain-of-function mutant protein with a premature stop (Q293*), which we find is freed from physical repression by its negative regulator, the PALS-22 protein. PALS-25(Q293*) activates two related gene expression programs, the Oomycete Recognition Response (ORR) against natural pathogens of the epidermis, and the Intracellular Pathogen Response (IPR) against natural intracellular pathogens of the intestine. A subset of ORR/IPR genes is upregulated in *pals-25(Q293*)* mutants, and they are resistant to oomycete infection in the epidermis, and microsporidia and virus infection in the intestine, but without compromising growth. Surprisingly, we find that activation of PALS-25 seems to primarily stimulate the downstream bZIP transcription factor ZIP-1 in the epidermis, which leads to upregulation of gene expression in both the epidermis and in the intestine. Interestingly, we find that this epidermal-to-intestinal signaling promotes resistance to the *N. parisii* intestinal pathogen, demonstrating cross-tissue protective immune induction from one epithelial tissue to another in *C. elegans*.

**Author summary:** Multicellular organisms need to monitor the health and function of multiple tissues simultaneously to respond appropriately to pathogen infection. Here, we study an ON/OFF switch in the roundworm *C. elegans* that controls immune responses to diverse natural pathogens of the skin and gut. We show a physical association between the ‘ON switch’ protein PALS-25 and the ‘OFF switch’ protein PALS-22, and that this association is disrupted in a mutant, activated form of PALS-25. When either PALS-22 is lost, or PALS-25 is activated, a downstream immune regulator ZIP-1 is activated specifically in the skin but not the gut. Excitingly, our findings show that skin-specific loss of PALS-22 or skin-specific activation of PALS-25 can induce immune responses in the worm gut. These findings highlight the coordination of immune responses across different tissues that are commonly infected by microbial pathogens.

## Introduction

How hosts coordinate defensive responses across different tissues during infection is an important question in immunology. Recent advances from a range of animal species have highlighted the importance of communication between the nervous system and peripheral tissues to regulate immunity, such as communication in the ‘brain-gut’ and ‘brain-skin’ axes [1–6]. However, the extent to which there is coordination of immune responses across peripheral tissues, such as a ‘skin-gut’ axis, is less well-understood [7].

The nematode *Caenorhabditis elegans* provides a convenient model to dissect whole animal immune responses to pathogen infection. *C. elegans* lacks known professional immune cells and instead seems to be highly dependent on epithelial cells of the epidermis and intestine for immune protection [8]. Immune signaling within these epithelial cells has been shown to be controlled by both proximal as well as distal signals. For example, the fungal pathogen *Drechmeria coniospora,* which infects the epidermis, causes upregulation of two classes of antimicrobial peptides, the neuropeptide-like protein (*nlp*) and the caenacin (*cnc*) genes. Upregulation of *nlp* genes is induced by a proximal cuticle-derived ligand, which triggers a G-protein protein-coupled receptor and subsequent PMK-1 p38 MAP kinase signaling in the epidermis [9, 10]. In contrast, infection-induced upregulation of the *cnc* genes is regulated more distally by the release of DBL-1 (a TGF-β analog) from neurons, leading to downstream TGF-β receptor signaling in epidermal cells, as part of a ‘brain-skin’ axis [11]. Similar themes have been found with immune responses to *Pseudomonas aeruginosa* and *Staphylococcus aureus*, bacterial pathogens that infect the intestinal lumen [2]. Infection with these pathogens induces expression in the intestine of a suite of host genes encoding predicted anti-microbials, and several studies have demonstrated that they can also be modulated cell non-autonomously by neurons as part of a ‘brain-gut’ axis [12–16].

Another example of immune regulation through the ‘brain-skin’ axis has recently been reported for response to the oomycete *Myzocytiopsis humicola,* which is a eukaryotic pathogen isolated from wild-caught *C. elegans* [17]. This pathogen infects through the *C. elegans* epidermis and induces upregulation of chitinase-like *(chil)* genes, which act to alter the properties of the cuticle and limit pathogen attachment [17]. Treatment of *C. elegans* with an innocuous extract derived from *M. humicola* is sufficient to induce *chil* gene expression in the epidermis, and extract also primes animals for enhanced pathogen resistance [18]. Interestingly, it was shown that upregulation of *chil* gene expression in the epidermis is dependent on neuronal TAX-2/TAX-4 signaling, suggesting that *M. humicola* extract is detected by neurons, which then signal to activate defense gene expression distally in the epidermis. Together, this host transcriptional response to infection by *M. humicola*, and to extract derived from *M. humicola,* comprise a novel immune program called the Oomycete Recognition Response (ORR) [18].

Interestingly, despite their shared epidermal tropism, and both being eukaryotic pathogens, the ORR induced by *M. humicola* infection has relatively little overlap with genes induced during *D. coniospora* infection [17]. Instead, the ORR shares more similarity with a host defense program that is upregulated in response to diverse intracellular pathogens of the intestine called the Intracellular Pathogen Response (IPR) [19, 20]. The IPR is induced following infection with the single-stranded RNA Orsay virus as well as the intracellular fungus *Nematocida parisii,* which belongs to the Microsporidia phylum [21–24]. Forward genetic screens revealed that both ORR and IPR gene expression are regulated by two members of the ‘<u>p</u>rotein containing <u>ALS2</u>cr12 signature’ *(pals)* gene family, which is expanded in *C. elegans* (39 members) compared to a single *pals* ortholog each in mice and humans [19, 25, 26]. While the *pals* genes have no known phenotypes in mammals, and *pals* genes encode no known biochemical function in any species, *pals-25* and *pals-22* cause important immune phenotypes in *C. elegans*, acting as an ON/OFF switch for the ORR and IPR programs. *pals-22* loss-of-function mutants have constitutive ORR and IPR gene expression, increased resistance to intracellular pathogens and *M. humicola*, increased tolerance of proteotoxic stress induced by heat shock (thermotolerance), increased RNA interference (RNAi), together with slowed growth and other phenotypes [18, 19, 26]. Loss-of-function mutations of *pals-25* suppress ORR and IPR gene expression and other mutant phenotypes described for *pals-22* mutants [19]. While *pals-22* acts in the intestine to regulate thermotolerance and some gene expression in the intestine, and *pals-22* can regulate immunity across a generation, it is not clear in which tissues either *pals-22* or *pals-25* acts to regulate immunity during infection [26, 27].

Here, we report a gain-of-function premature stop codon allele of *pals-25* (Q293*) that constitutively activates ORR and IPR gene expression in an otherwise wild-type background. Accordingly, we find that *pals-25(Q293*)* mutants share some, but not all phenotypes previously shown to be regulated by *pals-22/25*. In particular, we find that *pals-25(Q293*)* mutants have increased resistance to *M. humicola* infection in the epidermis, as well as Orsay virus and *N. parisii* infection in the intestine, but do not have increased thermotolerance or slowed growth. The *pals-25(Q293*)* mutation causes premature truncation of 13 amino acids in the C-terminus of the PALS-25 protein, which we find results in loss of physical interaction between PALS-25 and its repressor protein PALS-22. Surprisingly, we find that both loss of PALS-22 and activation of PALS-25 have stronger effects in the epidermis compared to the intestine when assessing nuclear localization of the bZIP transcription factor ZIP-1, which is required for inducing a subset of IPR genes [28]. Furthermore, we find that epidermal-specific expression of PALS-25(Q293*), or epidermal-specific loss of PALS-22 protein, induces IPR reporter expression in both the epidermis as well as the intestine, indicating cross-talk between these tissues. Finally, PALS-25 activity specifically in the epidermis was also able to promote resistance to intracellular pathogens of the intestine. Our results indicate that PALS-25 can signal cell non-autonomously across the ‘skin-gut’ axis to induce immune gene expression and promote systemic pathogen resistance across distinct peripheral tissues.

## Results

### A *pals-25* gain-of-function mutation activates pathogen response gene expression in the absence of infection

To identify regulators of the response to natural oomycete infection of the epidermis, we performed an EMS forward genetic screen (in the Barkoulas lab) for mutations that induce expression of the *chil-27p*::GFP reporter in the absence of infection. *chil-27* is upregulated in response to *M. humicola* oomycete infection of the epidermis as part of the ORR [17, 18]. Here, an F2 screen identified the *icb98* mutation, which leads to constitutive *chil-27p*::GFP expression in the epidermis. Whole-genome sequencing revealed that *icb98* is a mutation in the *pals-25* gene, predicted to convert amino acid Q293 into a premature stop codon (Q293*), causing the loss of the C-terminal 13 amino acids of PALS-25 protein (Fig 1A, B S1 and S2 Tables). Because previous studies indicated that wild-type PALS-25 was an activator of *chil-27* expression [19], we hypothesized that *icb98* was a gain-of-function mutation. To test this hypothesis, and to determine whether the predicted Q293* mutation in PALS-25 protein was the cause of the *icb98* mutant phenotype, we performed RNAi against *pals-25* and found that constitutive expression of the *chil-27p*::GFP reporter was suppressed (Fig 1C). This result indicated that *icb98* is a *pals-25* gain-of-function allele, which is responsible for constitutive activation of *chil-27p*::GFP expression.

**Fig 1.**
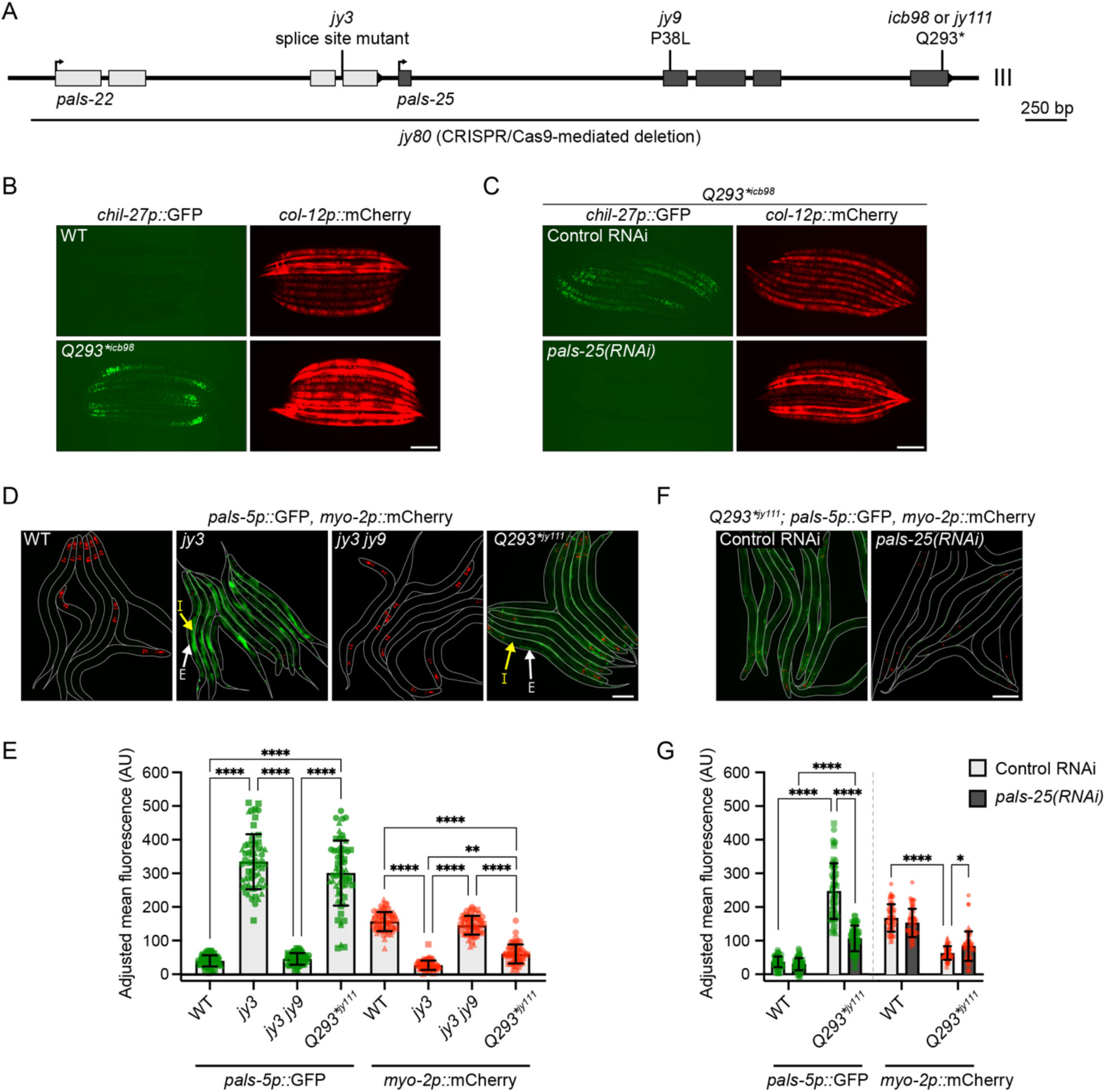
A gain-of-function allele of *pals-25* activates immune responses in the absence of infection. **A)** The *pals-22* and *pals-25* gene coding structure, which comprises an operon. *pals-22* exons are shown in light grey boxes and *pals-25* exons are dark grey, UTR regions not shown. Positions of mutant alleles in this study are indicated and residues altered are described in S2 Table. Due to space constraints, *pals-22(jy3)* and *pals-25(jy9, jy111* and *icb98)* allele names will be used without gene names in this and subsequent figures. *icb98* and *jy111* are independently isolated *pals-25* gain-of-function alleles with the same *Q293\** change, and will be designated in figures as *Q293*^icb98^* and *Q293*^jy111^*, respectively. **B)** *pals-25(Q293*)^icb98^* mutants show upregulated expression of *chil-27p::*GFP compared to WT, in the absence of infection. *col-12p::mCherry* is part of the same transgene and is constitutively expressed in both WT and *pals-25(Q293*)^icb98^*. L4 and young adult animals are shown. **C)** Expression of *chil-27p::*GFP in *pals-25(Q293*)^icb98^* is lost upon treatment with *pals-25(RNAi)*. L4 and young adult animals are shown. **D)** *pals-22(jy3)* and *pals-25(Q293*)^jy111^* mutants show constitutive expression of *pals-5p::*GFP. *myo-2p::mCherry* is part of the same transgene and is constitutively expressed in the pharynx of transgenic animals at all life stages. *pals-22(jy3)* mutants show *pals-5p::*GFP expression in both the intestine and epidermis while *pals-25(Q293*)^jy111^* mutants show expression more prominently in the epidermis. Yellow arrows indicate intestinal tissue (I) and white arrows indicate epidermal tissue (E). L4s are shown for all genotypes, with lines denoting the outline of the body. **E)** Quantification of *pals-5p::*GFP and *myo-2p::*mCherry mean fluorescence signal normalized to body area at L4. *pals-22(jy3)* and *pals-25(Q293*)^jy111^* mutants exhibit both *pals-5p::*GFP induction, and transgene silencing as measured by *myo-2p::*mCherry fluorescence, which is significantly different from wild type (WT). Kruskal-Wallis test with Dunn’s multiple comparisons test with each reporter analyzed independently. **F)** Expression of *pals-5p::*GFP in *pals-25(Q293*)^jy111^* mutants is suppressed upon treatment with *pals-25(RNAi)*. L4s are shown, with lines denoting the outline of the body. **G)** Quantification of *pals-5p::*GFP and *myo-2p::*mCherry mean fluorescence signal normalized to body area at L4 for different RNAi treatments. Two-way ANOVA with Sidak’s multiple comparisons test with each reporter analyzed independently. For **E** and **G**, n = 60 animals per genotype or treatment, three experimental replicates. Symbols represent fluorescence measurements for individual animals and different symbol shapes represent animals from experimental replicates performed on different days. Bar heights indicate mean values and error bars represent standard deviations. **** *p* < 0.0001, ** *p* < 0.01, * *p* < 0.05. For **B-D** and **F**, scale bar = 100 µm.

In addition to regulating expression of ORR genes, *pals-25* had previously been shown to regulate expression of IPR genes, which comprise an overlapping but distinct gene set from the ORR gene set [18, 19]. To investigate whether the predicted Q293* mutation of the PALS-25 protein would also induce IPR gene expression in a wild-type background, we used CRISPR/Cas9-mediated mutagenesis (in the Troemel lab) to independently generate this change in a separate strain background carrying the *pals-5p::*GFP IPR reporter [29]. Indeed, the resulting *pals-25(Q293*)^jy111^* allele induced constitutive expression of *pals-5p::*GFP, like the previously described phenotype of the *jy3* loss-of-function mutation in *pals-22* (Fig 1A, D-E, S1 and S2 Tables) [26]. Of note, *pals-22(jy3)* mutants displayed strong *pals-5p::*GFP expression in the epidermis, pharynx, neurons, and intestine, while *pals-25(Q293*)^jy111^* mutants displayed *pals-5p::*GFP expression in similar tissues, but with weaker expression in the intestine (Fig 1D). Again, consistent with *pals-25(Q293*)^jy111^* being a gain-of-function mutation, we found that RNAi knockdown of *pals-25* in *pals-25(Q293*)^jy111^* mutants suppressed constitutive expression of the *pals-5p::*GFP reporter (Fig 1F, G).

Besides their roles in regulating pathogen response gene expression, *pals-22* and *pals-25* are also known to regulate transgene silencing, with *pals-22* mutants showing increased transgene silencing that is suppressed by *pals-25* mutations [19, 25, 26]. Therefore, we analyzed transgene silencing by measuring fluorescence intensity of the *myo-2p::*mCherry co-injection marker that is co-expressed with the *pals-5p::*GFP reporter as part of the same *jyIs8* transgene, but is not regulated by infection. Here we found that *pals-25(Q293*)^jy111^* suppressed expression of *myo-2p*::mCherry, indicating that like *pals-22(jy3)* loss-of-function mutants, the *pals-25(Q293*)^jy111^* mutants have increased transgene silencing, which is suppressed by *pals-25* RNAi (Fig 1D-G). In summary, *pals-25(Q293*)* mutants have increased expression of the ORR and IPR reporters, as well as increased transgene silencing. All of these phenotypes can be suppressed by RNAi against *pals-25* (Fig 1C, F, G), suggesting that the predicted Q293* change in PALS-25 leads to activation of this protein in a wild-type background.

### A subset of ORR and IPR genes are upregulated in *pals-25(Q293*)* mutants

Given the ability of *pals-25(Q293*)* mutations to induce *chil-27p::*GFP and *pals-5p::*GFP reporter expression, we next investigated endogenous mRNA expression of ORR and IPR genes in these mutants. First, we performed qRT-PCR to analyze endogenous mRNA expression of *pals-5*, *chil-27*, and other known ORR and IPR genes in *pals-25(Q293*)^jy111^* mutants along with wild-type animals, *pals-22(jy3)* mutants, and *pals-22(jy3) pals-25(jy9)* double mutants as controls (S3 Table). As a positive control, we found that *pals-22(jy3)* mutants displayed constitutive upregulation of *pals-5* (an ORR/IPR gene), *chil-27* (an ORR gene), and other ORR and IPR genes when compared to wild-type, and induction of these genes was suppressed in *pals-22(jy3) pals-25(jy9)* double mutants (Fig 2A). In contrast, while *pals-25(Q293*)^jy111^* mutants displayed upregulation of *pals-5, chil-27,* and some ORR/IPR genes, expression levels of ORR/IPR genes were not always upregulated to the same degree as observed in *pals-22(jy3)* mutants, or in the case of F26F2.1 (an IPR gene), no upregulation relative to wild-type animals was observed (Fig 2A). Thus, *pals-25(Q293*)^jy111^* mutants appear to upregulate some, but not all of the genes induced by wild-type *pals-25* in a *pals-22* mutant background.

**Fig 2.**
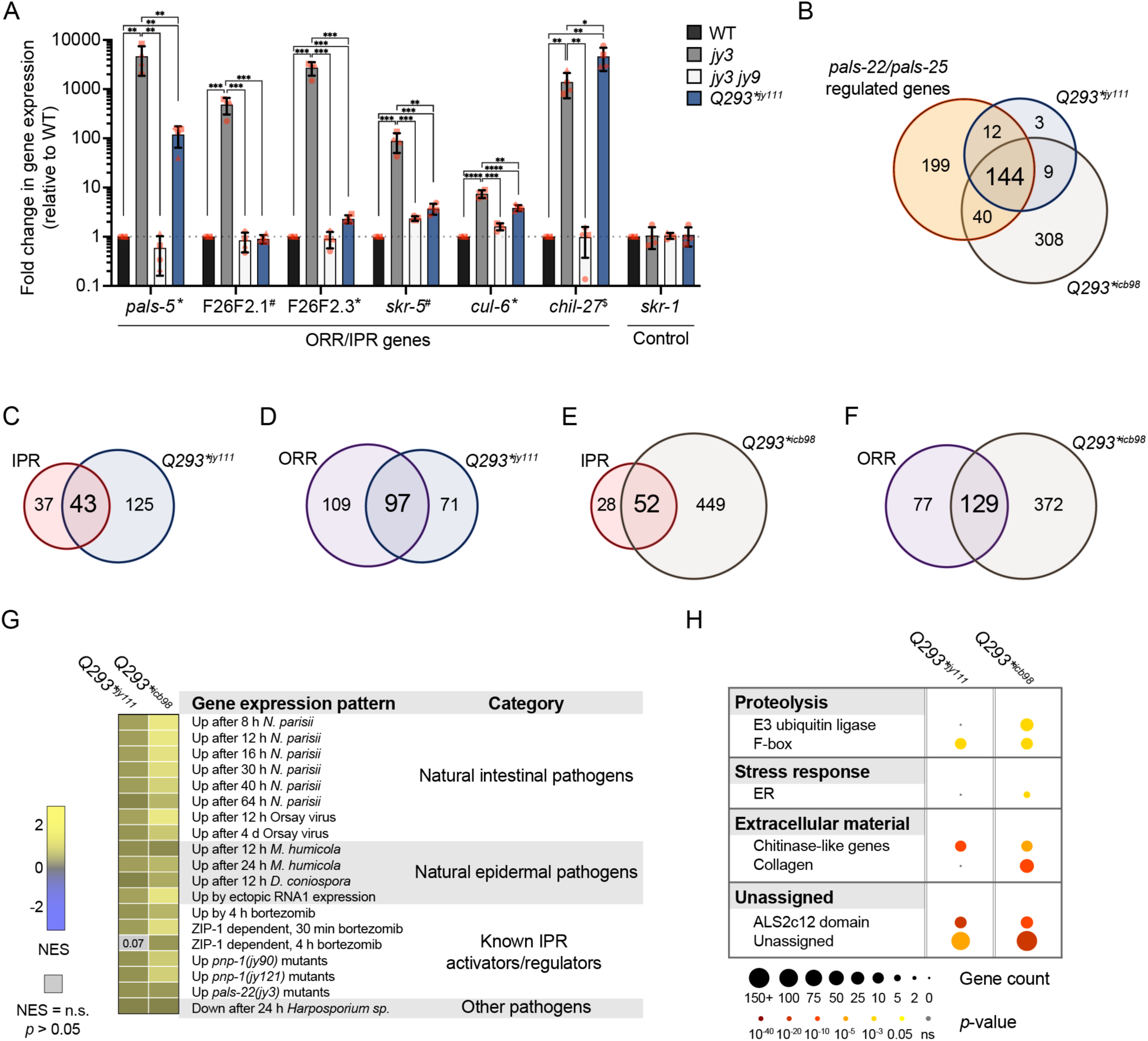
A subset of immune response genes is upregulated in *pals-25(Q293*)* mutants. **A)** qRT-PCR of select ORR and IPR genes. * = Gene belongs to ORR and IPR, # = Gene belongs to only IPR, $ = Gene belongs to only ORR. The results shown are fold change in gene expression relative to WT. **** *p* < 0.0001, *** *p* < 0.001, ** *p* < 0.01, * *p* < 0.05, One-tailed t-test. n = 4 independent experimental replicates, different symbol shapes represent the expression values for replicates performed on different days. Bar heights indicate mean values and error bars represent standard deviations. **B)** Upregulated genes in both *pals-25(Q293*)^jy111^* and *pals-25(Q293*)^icb98^* mutants have significant overlap with genes regulated by wild-type *pals-22* and *pals-25*. Hypergeometric test, RF = 34.5; *p* < 5.39e-243 and RF = 15.6; *p* < 1.994e-182 for *pals-25(Q293*)^jy111^* and *pals-25(Q293*)^icb98^*, respectively. **C-D)** Upregulated genes in *pals-25(Q293*)^jy111^* mutants have significant overlap with the IPR and ORR. Hypergeometric test, IPR: RF = 46.9; *p* < 6.215e-64, ORR: RF = 41.1; *p* < 6.883e-144. **E-F)** Upregulated genes in *pals-25(Q293*)^icb98^* mutants have significant overlap with the IPR and ORR. Hypergeometric test, IPR: RF = 21.8; *p* < 4.529e-60, ORR: RF = 21; *p* < 5.122e-148. **G)** Correlation of differentially expressed genes in *pals-25(Q293*)^jy111^* and *pals-25(Q293*)^icb98^* mutants with those expressed during pathogen infection and regulated by known activators of the IPR. Correlation of gene sets quantified as Normalized Enrichment Score (NES). Blue indicates significant correlation of downregulated genes in *pals-25(Q293*)^jy111^* or *pals-25(Q293*)^icb98^*mutants with the gene sets tested, and yellow indicates significant correlation of upregulated genes in *pals-25(Q293*)^jy111^* or *pals-25(Q293*)^icb98^* mutants with the gene sets tested. Grey indicates no significant correlation (*p* > 0.05 or False Discovery Rate > 0.25). GSEA analysis of 93 gene sets tested in S7 Table. **H)** WormCat analysis shows significantly enriched gene categories represented in upregulated genes of *pals-25(Q293*)^jy111^* and *pals-25(Q293*)^icb98^* mutants. Bold text indicates broad “Category 1” biological processes enriched while nested text indicates more specific “Category 2 or 3” processes enriched. *p* values were determined using Fisher’s exact test with Bonferroni correction from minimum hypergeometric scores calculated in the WormCat software. A summary of WormCat analysis can be found in S8 Table.

To more broadly investigate ORR and IPR gene expression induced by *pals-25(Q293*)* mutations, we performed independent RNAseq analysis of *pals-25(Q293*)^jy111^* mutants (in the Troemel lab) and *pals-25(Q293*)^icb98^* mutants (in the Barkoulas lab). Differential gene expression analysis determined that 168 genes are upregulated in *pals-25(Q293*)^jy111^* mutants (*p* < 0.05, log_2_ fold change > 2) and 501 genes are upregulated in *pals-25(Q293*)^icb98^* mutants (*p* < 0.01, log_2_ fold change > 2) compared to wild-type animals (S4 and S5 Tables). Genes upregulated in *pals-25(Q293*)^jy111^* and *pals-25(Q293*)^icb98^* mutants overlap significantly with each other (153/168 genes up *in Q293*^jy111^* are also up in *Q293*^icb98^*), highlighting the robustness of these results for *pals-25(Q293*)* mutants, performed in different strain backgrounds and different laboratories. *pals-25(Q293*)* upregulated genes also overlap significantly with those regulated by wild-type *pals-22/25*, i.e. genes that are upregulated both in *pals-22(jy3)* mutants when compared to wild-type animals, and in *pals-22(jy3)* mutants when compared to *pals-22(jy3) pals-25(jy9)* double mutants (S1 Fig, S6 Table). However, the overlap is only partial, with less than half of the *pals-22/25*-regulated genes also upregulated by *pals-25(Q293*)* (Fig 2B, S6 Table). Similarly, there was partial overlap with ORR and IPR genes, which are generally a subset of *pals-22/25*-regulated genes; here we saw that approximately one-half to two-thirds of the IPR and ORR genes were upregulated in *pals-25(Q293*)* mutants (Fig 2C-F, S2 Fig, S6 Table).

To compare the gene sets altered in our two *pals-25(Q293*)* mutant strains to gene sets regulated in response to infection by a variety of pathogens, previously characterized IPR activators, exogenous stresses, and other known immunity regulators, we performed Gene Set Enrichment Analysis (GSEA) [30]. Here we found that both *pals-25(Q293*)^jy111^* and *pals-25(Q293*)^icb98^* mutants upregulate genes in common with natural infections by intestinal and epidermal pathogens as well as known activators and regulators of the IPR (Fig 2G, S7 Table). To determine categories of genes that are upregulated in *pals-25(Q293*)^jy111^* and *pals-25(Q293*)^icb98^* mutants we performed WormCat analysis [31]. Here we found genes involved in proteolysis including E3 ubiquitin ligase and F-box genes, ER stress response genes, genes involved in the regulation of extracellular material such as collagen and chitinase-like genes, and notably genes with a *pals* domain (Fig 2H, S8 Table). The categories of genes represented in *pals-25(Q293*)^jy111^* and *pals-25(Q293*)^icb98^* mutants are similar to those previously reported to be represented in *pals-22(jy3)* mutants, although *pals-22(jy3)* mutants show additional stress response genes related to pathogen infection that are not represented in *pals-25(Q293*)* mutants [32]. Together, our analysis demonstrates that a subset of genes previously defined as being induced by wild-type *pals-25* (i.e. genes induced in *pals-22* mutants in a *pals-25*-dependent manner) are also upregulated in *pals-25(Q293*)* mutants, including genes induced by natural pathogen infection.

### *pals-25(Q293*)* mutants have increased immunity against natural pathogens, but do not have other phenotypes associated with the IPR

Next, we investigated *pals-25(Q293*)* mutants for phenotypes associated with ORR/IPR activation, i.e. phenotypes found in *pals-22* mutants that depend on wild-type *pals-25*. Because *pals-25(Q293*)^icb98^* was isolated in the screen for ORR regulation, we first investigated oomycete resistance in *pals-25(Q293*)^icb98^* mutants and found that they have increased survival upon *M. humicola* infection when compared to wild-type animals (Fig 3A). *Because pals-25(Q293*)^jy111^* was shown to regulate IPR expression, we next analyzed *pals-25(Q293*)^jy111^* resistance to *N. parisii.* Here we quantified sporoplasms [33], which are mononucleate intracellular parasite cells and found that *pals-25(Q293*)^jy111^* mutants are resistant to *N. parisii* at 3 hours post-infection (hpi) relative to wild-type animals, and the degree of resistance was indistinguishable from *pals-22(jy3)* mutants (Fig 3B). We also analyzed *pals-25(Q293*)^jy111^* resistance at 30 hpi, when there is active *N. parisii* replication as multinucleate meront cells inside of host intestinal cells [34]. Here again, we found that *pals-25(Q293*)^jy111^* mutants, similar to *pals-22(jy3)* mutants, are resistant to *N. parisii* compared to wild-type animals (Fig 3C, S3A Fig). Finally, we analyzed *pals-25(Q293*)^jy111^* mutant resistance to Orsay virus at 18 hpi. Here, we found that *pals-25(Q293*)^jy111^* mutants are resistant to viral infection when compared to wild-type animals, although not quite as resistant as *pals-22(jy3)* mutants (Fig 3D, S3B Fig). Fluorescence bead-feeding control assays indicate that *pals-25(Q293*)^jy111^* mutants do not have a feeding impairment that would reduce uptake and exposure to pathogens in the intestinal lumen, but instead they appear to have bona fide increased immunity against intracellular intestinal pathogens (S4 Fig).

**Fig 3.**
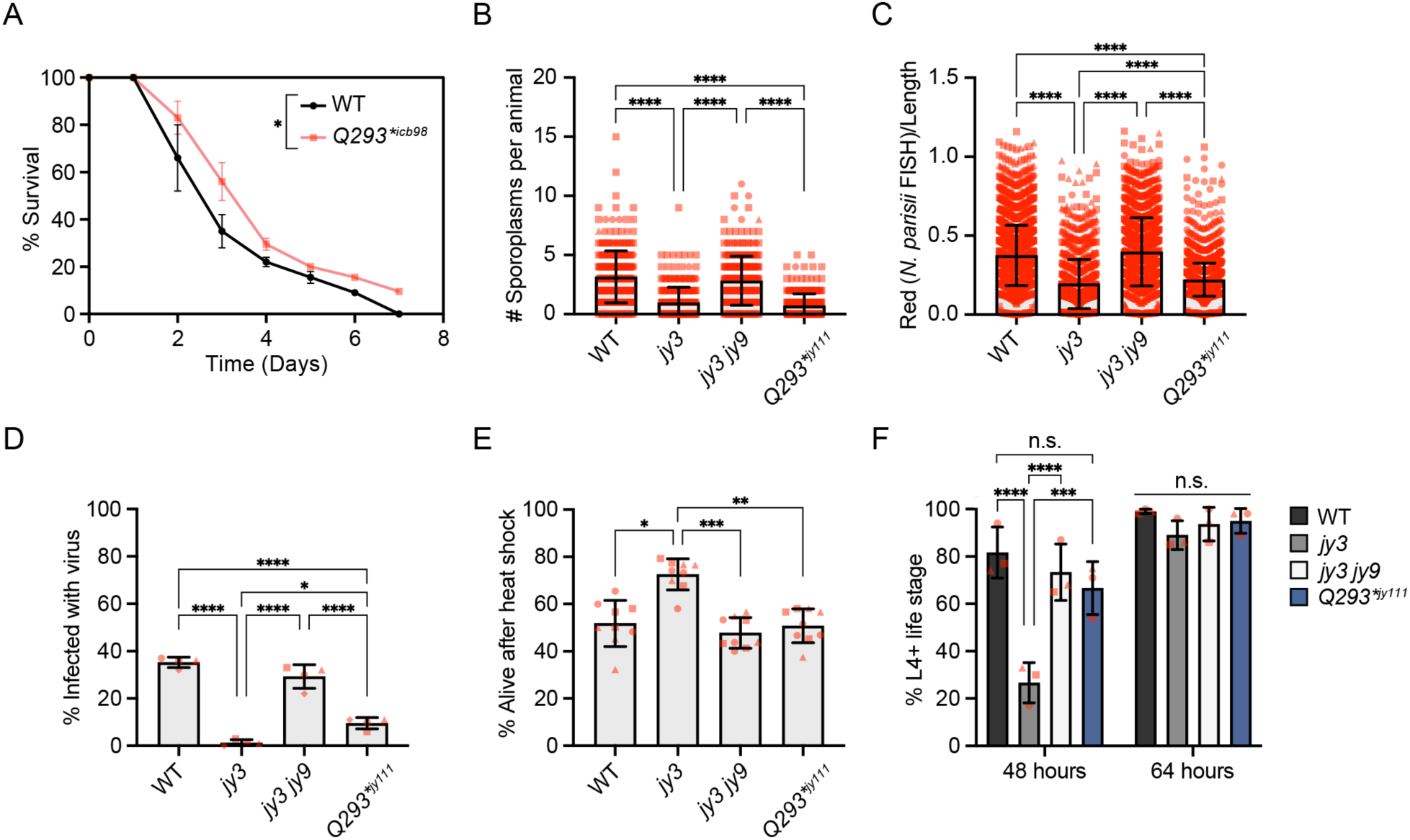
*pals-25(Q293*)* mutants have increased resistance to natural pathogens but not increased resistance to proteotoxic stress, nor developmental delay. **A)** *pals-25(Q293*)^icb98^* mutants have increased survival upon infection with *M. humicola* as compared to WT animals. Log-rank test. n = 90 animals across three experimental replicates. **B-C)** *pals-22(jy3)* and *pals-25(Q293*)^jy111^* mutants exhibit resistance to *N. parisii* compared to WT at 3 hours post infection (hpi) **(B)** and 30 hpi **(C)**. Kruskal-Wallis test with Dunn’s multiple comparisons test. 3 hpi: n = 300 animals per genotype, three experimental replicates. Symbols represent the number of *N. parisii* sporoplasms infecting an individual animal, as determined by *N. parisii*-specific fluorescent FISH signal. 30 hpi: n = WT: 2,972, *pals-22(jy3)*: 1,151, *pals-22(jy3) pals-25(jy9)*: 1,837, *pals-25(Q293*)^jy111^* : 3,204 animals, three experimental replicates. Symbols represent the infected area of individual worms by FISH fluorescent signal normalized to time-of-flight quantified on a COPAS Biosort machine. **D)** *pals-22(jy3)* and *pals-25(Q293*)^jy111^* mutants exhibit resistance to Orsay virus compared to WT at 18 hpi. One-way ANOVA with Tukey’s multiple comparisons test. n = 4 experimental replicates, 100 animals scored in each replicate per genotype. **E)** *pals-22(jy3)* mutants exhibit increased survival compared to WT after 2 h of heat shock at 37.5 °C, followed by 24 h at 20 °C, but *pals-25(Q293*)^jy111^* mutants do not. Kruskal-Wallis test with Dunn’s multiple comparisons test. n = 9 plates, 30 animals per plate, tested in triplicate. **F)** *pals-22(jy3)* mutants, but not *pals-25(Q293*)^jy111^* mutants, exhibit developmental delay compared to WT. Two-way ANOVA with Sidak’s multiple comparisons test for simple effects within a timepoint. For **A-F**, **** *p* < 0.0001, *** *p* < 0.001, ** *p* < 0.01, * *p* < 0.05. For **B-F**, bar heights indicate mean values and error bars represent standard deviations. Different symbol shapes represent data points from assays performed on different days.

Additional phenotypes found in *pals-22* mutants that are dependent upon wild-type *pals-25* include increased thermotolerance and slowed development relative to wild-type animals [19]. To determine if *pals-25(Q293*)^jy111^* mutants have increased thermotolerance we subjected them to heat shock (37.5 °C for 2 h), and measured survival after 24 h at 20 °C. Consistent with previous results, we found that *pals-22(jy3)* mutants display increased thermotolerance in this assay, which is suppressed to wild-type levels in *pals-22(jy3) pals-25(jy9)* double mutants (Fig 3E). However, we found that *pals-25(Q293*)^jy111^* mutant survival was indistinguishable from wild-type animals 24 h after heat shock (Fig 3E), indicating they do not have increased thermotolerance. We also assessed the developmental rate of *pals-25(Q293*)^jy111^* mutants by monitoring the time for synchronized embryos to reach the L4 life stage or older when grown at 20 °C. Similar to previous results, at 48 h we found that >80% of wild-type animals had reached the L4 stage while <30% of *pals-22(jy3)* mutants had reached L4, and this phenotype was significantly suppressed by a *pals-25(jy9)* mutation. In contrast, almost 70% of *pals-25(Q293*)^jy111^* mutants had reached the L4 stage, similar to wild-type animals (Fig 3F). Taken together, our results demonstrate that *pals-25(Q293*)* mutants do not share all the phenotypes exhibited by *pals-22* mutants, but they do display resistance to natural pathogens and increased transgene silencing similar to *pals-22* mutants. All of these phenotypes had previously been shown to be regulated by *pals-25* loss-of-function mutations only in a *pals-22* mutant background [19], while the results described above demonstrate that *pals-25(Q293*)* mutants can specifically regulate infection-related phenotypes in a wild-type background.

### The truncated PALS-25(Q293*) protein does not associate with the PALS-22 protein

One possible explanation for why *pals-25(Q293*)* mutants display ORR and IPR phenotypes in a wild-type background is that the PALS-25(Q293*) protein is somehow freed from repression by the wild-type PALS-22 protein. Previous co-immunoprecipitation and mass spectrometry studies (co-IP/MS) suggested that PALS-22 and PALS-25 are physically associated, leading to the hypothesis that PALS-25 is repressed by PALS-22 through this physical interaction [35]. To analyze PALS-22/PALS-25 interactions, we performed co-IP of PALS-22 using strains where PALS-22 is C-terminally tagged with GFP::3xFLAG, and then performed Western blots using antibodies we raised against endogenous PALS-25 protein to determine if wild-type (WT) PALS-25 would be recovered by co-IP with PALS-22. Here, we found that PALS-25(WT) was recovered in co-IP experiments performed on a strain where PALS-22 was expressed in single-copy from the MosSCI locus [26, 36] with its endogenous promoter (*pals-22p*), confirming previous co-IP/MS results showing a physical association between PALS-22 and PALS-25(WT) proteins (Fig 4A).

**Fig 4.**
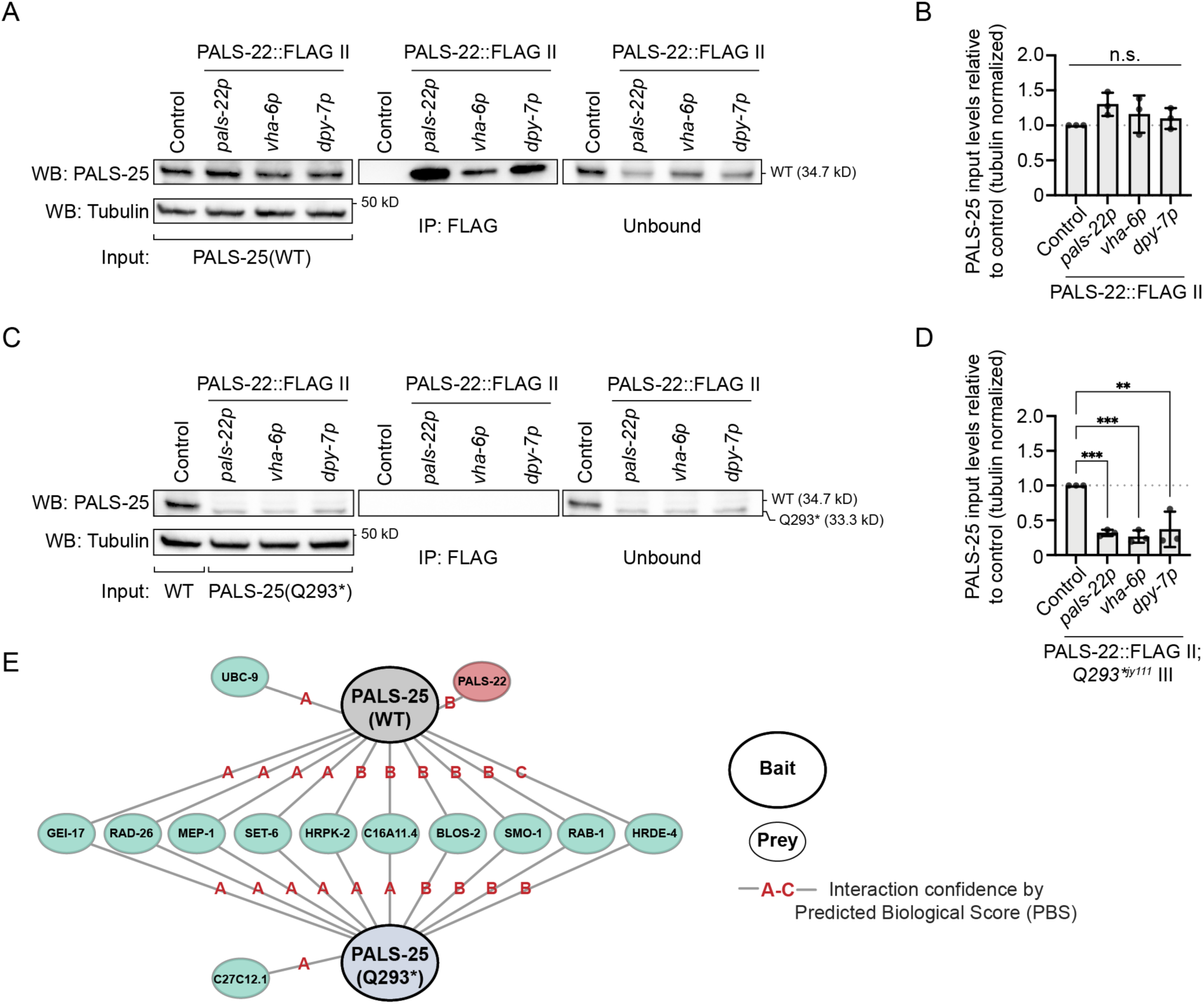
PALS-25(Q293*) protein is not physically associated with PALS-22 protein. **A)** Co-immunoprecipitation (co-IP) of FLAG-tagged PALS-22 and Western blot (WB) analysis for PALS-25(WT) in different tissues. PALS-22::GFP::3xFLAG expressed from its endogenous promoter (*pals-22p*), intestinal promoter (*vha-6p*), or epidermal promoter (*dpy-7p*) captured by FLAG-IP interacts with endogenously expressed PALS-25(WT). A GFP::3xFLAG control expressed from the *spp-5p* promoter does not interact with PALS-25(WT). **B)** Quantification of PALS-25 input levels relative to the control strain. PALS-25 input levels are similar for the strains used in **A** for co-IP/WB. *p* > 0.05, one-way ANOVA with Dunnett’s multiple comparisons test with respect to the GFP::FLAG control strain. n = 3 independent co-IP/WB experimental replicates. **C)** co-IP of FLAG-tagged PALS-22 expressed in different tissues, as described in **A**, and WB analysis for PALS-25(Q293*). PALS-22 captured by FLAG-IP does not interact with endogenously expressed PALS-25(Q293*). The GFP::3xFLAG control does not interact with PALS-25(Q293*). **D)** Quantification of PALS-25(Q293*) input levels relative to PALS-25(WT) in the control strain. PALS-25(Q293*) protein levels are decreased relative to control for the strains used in **C** for co-IP/WB. *** *p* < 0.001, ** *p* < 0.01, one-way ANOVA with Dunnett’s multiple comparisons test to GFP::FLAG control strain. n = 3 independent co-IP/WB experimental replicates. **E)** Yeast two-hybrid analysis of PALS-25(WT) and PALS-25(Q293*) baits with interacting prey proteins from a proteome-wide screen. PALS-25(WT) interacts with PALS-22 but PALS-25(Q293*) does not. Lines indicate significant interaction between bait and prey. Red letters indicate confidence of interaction from A (highest) to C (lowest) as determined by Predicted Biological Score (PBS). A detailed description of PBS analysis and complete bait-prey interactions is listed in the Materials and Methods and S10 Table.

We next analyzed in which tissues PALS-22 and PALS-25(WT) proteins interact. Based on fosmid transgene analysis [37], PALS-22 and PALS-25(WT) proteins are expressed in several tissues, including the epidermis and intestine (S5 Fig, S9 Table). To analyze PALS-22 and PALS-25 protein-protein interaction in a tissue-specific way, we used strains where PALS-22::GFP::3xFLAG was expressed in the intestine with an intestinal-specific promoter (*vha-6p*), or in the epidermis with an epidermal-specific promoter (*dpy-7p*) from the MosSCI locus [26, 36]. Here we found that PALS-22 and PALS-25(WT) proteins were associated with each other in both the intestine and in the epidermis (Fig 4A, B).

We next asked if the association between PALS-22 and PALS-25 might be lost for the C-terminally truncated PALS-25(Q293*) mutants. Importantly, Western blots of whole *C. elegans* lysate revealed a lower molecular weight band for PALS-25(Q293*) protein, consistent with having lost the C terminal 13 amino acids (Fig 4C). Interestingly, unlike with PALS-25(WT), we did not observe co-IP of PALS-25(Q293*) with PALS-22 protein in any of the strains and tissues we tested (Fig 4C), suggesting that PALS-25(Q293*) no longer associates with PALS-22. The absence of PALS-25(Q293*) co-IP was not due to failure to capture PALS-22::GFP::3xFLAG, as PALS-22 was successfully immunoprecipitated in all samples (S6 Fig). However, we noted that, relative to PALS-25(WT) protein, expression of the PALS-25(Q293*) protein was significantly lower, despite using the same concentration of total protein extract for all samples (Fig 4A-D and Materials and Methods). Therefore, the lack of PALS-25(Q293*) co-IP with PALS-22 could be due to these lowered PALS-25 protein levels. To control for this difference, we performed a yeast two-hybrid screen as a complement to our co-IP/Western blot analysis, screening either PALS-25(WT) or PALS-25(Q293*) baits against a proteome-wide library of *C. elegans* prey proteins. Here we found that PALS-25(WT) and PALS-25(Q293*) interacted with almost all of the same proteins, except that PALS-22 was a high confidence hit for interaction only with PALS-25(WT), and not with PALS-25(Q293*) (Fig 4E, S10 Table). Thus, using two distinct biological systems and assays, we found that PALS-25(Q293*) protein loses its normal physical association with PALS-22 protein, providing a likely explanation for how PALS-25(Q293*) is free from repression by PALS-22 protein, and can activate the ORR, IPR, and pathogen resistance phenotypes in a background where *pals-22* is wild-type.

### The transcription factor ZIP-1 is activated downstream of PALS-25(Q293*) in the epidermis

Next, we investigated whether the bZIP transcription factor ZIP-1 is required downstream of PALS-25(Q293*) to activate pathogen response gene expression. ZIP-1 is required to induce the expression of some IPR genes in response to multiple triggers, including loss of *pals-22* [28]. Here, we found that *zip-1(RNAi)* significantly decreased *pals-5*p::GFP expression in *pals-25(Q293*)^jy111^* mutants relative to control RNAi, while the transgene silencing phenotype of *myo-2p::*mCherry observed in *pals-25(Q293*)^jy111^* mutants was unaffected by *zip-1(RNAi)* (Fig 5A, B). Therefore, ZIP-1 is required for the constitutive *pals-5p*::GFP gene expression in *pals-25(Q293*)^jy111^* mutants, but ZIP-1 does not appear to be required for transgene silencing.

**Fig 5.**
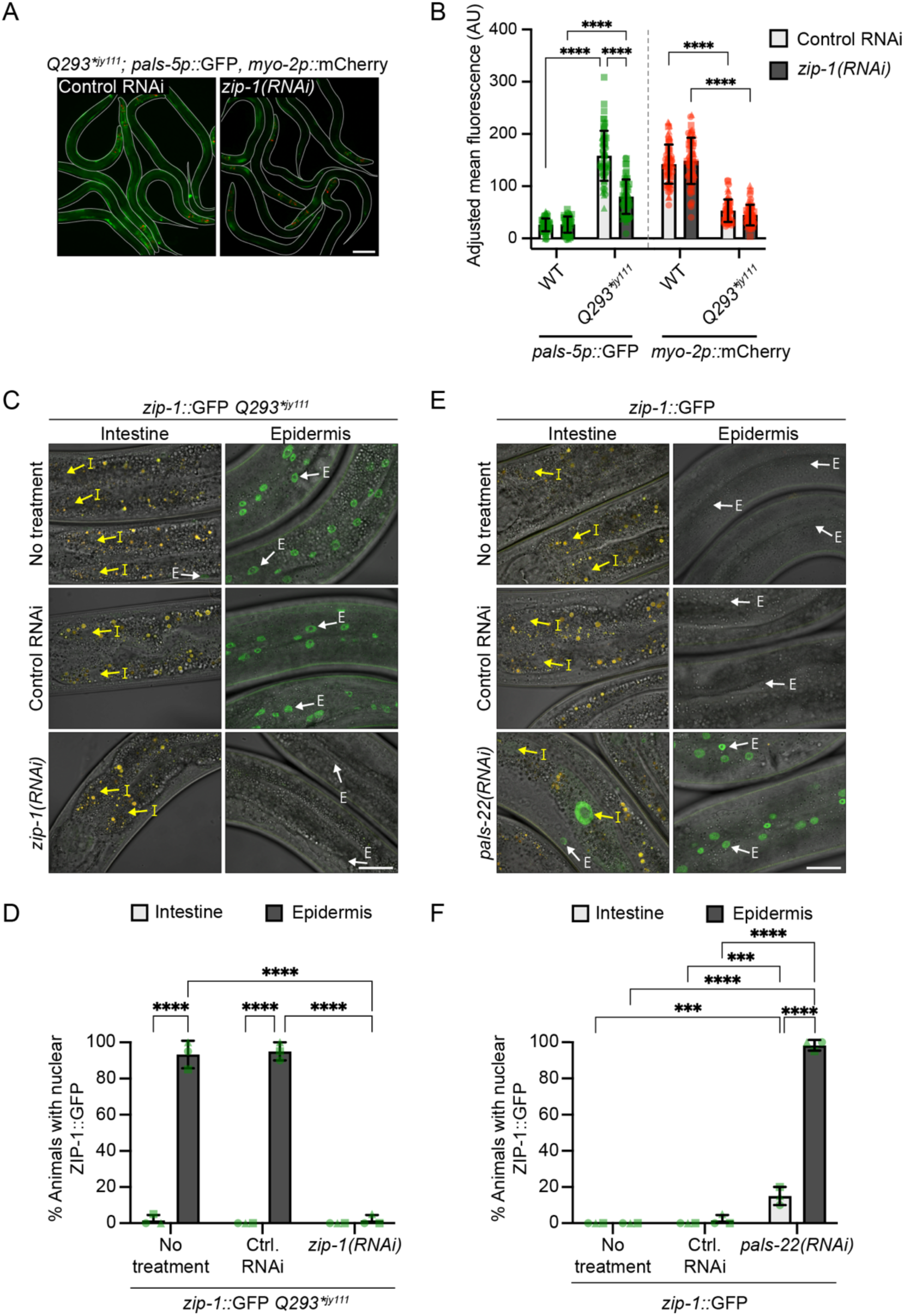
PALS-25(Q293*) and loss of PALS-22 protein both increase ZIP-1 nuclear localization in the epidermis. **A)** Constitutive expression of *pals-5p::*GFP in *pals-25(Q293*)^jy111^* mutants is suppressed upon treatment with *zip-1(RNAi)*. L4s are shown, with lines denoting the outline of the body. Scale bar = 100 µm. **B)** Quantification of *pals-5p::*GFP and *myo-2p::*mCherry mean fluorescence signal normalized to body area for RNAi-treated L4s. Symbols represent fluorescence measurements for individual animals, n = 60 animals per treatment. **C, D)** ZIP-1::GFP is constitutively expressed in the epidermis but not the intestine of *pals-25(Q293*)^jy111^* animals. **E, F)** ZIP-1::GFP is expressed in epidermal nuclei following *pals-22(RNAi)* but rarely in intestinal nuclei, and almost never in untreated or control RNAi-treated worms. For **C** and **E**, yellow arrows highlight intestinal nuclei ‘I’ and white arrows highlight epidermal nuclei ‘E’. Images are a composite of differential interference contrast, GFP, and RFP fluorescence channels and auto-fluorescent intestinal gut granules appear as yellow signal. Scale bar = 20 µm. For **B, D** and **F** bar heights indicate mean values and error bars represent standard deviations. **** *p* < 0.0001, *** *p* < 0.001, two-way ANOVA with Sidak’s multiple comparisons test (for **B,** each reporter analyzed independently). For **D** and **F**, n = 3 experimental replicates, 20 animals per treatment assessed for both intestinal and epidermal ZIP-1::GFP expression for each replicate. Different symbols represent replicates performed on different days.

Previous studies indicated that ZIP-1 localizes to the nucleus in response to various IPR triggers, including the proteasome-inhibitor bortezomib [28]. Indeed, we found that bortezomib induced ZIP-1::GFP nuclear expression in intestinal nuclei in 97% (58/60) of BTZ-treated animals compared to 0% (0/60) of control animals (S7 Fig). BTZ also induced ZIP-1::GFP localization in epidermal nuclei, but in only 30% (18/60) of animals (S7 Fig). We next examined nuclear localization of ZIP-1::GFP in *pals-25(Q293*)^jy111^* mutants. In contrast to BTZ treatment, we found strong ZIP-1::GFP nuclear localization in epidermal nuclei of *pals-25(Q293*)^jy111^* mutants, with ∼93% (56/60) of animals displaying nuclear localization (Fig 5C, D). Interestingly, we rarely observed intestinal nuclear localization of ZIP-1::GFP in *zip-1::gfp pals-25(Q293*)^jy111^* animals (Fig 5C, D). Thus, PALS-25(Q293*) appears to strongly activate ZIP-1::GFP in epidermal cells, but not in intestinal cells.

To determine if epidermal-biased nuclear localization of ZIP-1 was specific to *pals-25(Q293*)*, or is a more general feature of PALS-25 activation, we performed *pals-22(RNAi)* on *zip-1::gfp* animals and examined ZIP-1::GFP localization in intestinal and epidermal tissues. Similar to what we observed with *pals-25(Q293*)^jy111^* mutants, we found >98% (59/60) of *pals-22(RNAi)*-treated animals had nuclear localization of ZIP-1 in the epidermis, with very rare nuclear expression in the epidermis of control animals (Fig 5E, F). Also similar to *pals-25(Q293*)^jy111^* mutants, we observed only a low level of nuclear localization of ZIP-1::GFP in the intestine after *pals-22(RNAi)* (9/60, 15%), and never in control conditions (0/60) (Fig 5E, F). Thus, our results suggest that in contrast to BTZ treatment, which regulates ZIP-1 nuclear localization predominantly in intestinal cells, PALS-22 and PALS-25 regulate activation of the transcription factor ZIP-1 almost exclusively in epidermal cells.

### Epidermis-specific activation of PALS-25 signals to the intestine to promote pathogen resistance

Our results above suggest that PALS-22 and PALS-25 regulate the transcription factor ZIP-1 mostly in the epidermis (Fig 5). Furthermore, PALS-22 and PALS-25 proteins are both strongly expressed in epidermal tissue (S5 Fig). Therefore, we investigated whether activation of PALS-25 only in the epidermis might be sufficient to promote an immune response in both the epidermis and intestine. To test this hypothesis, we first developed a construct where *pals-25(Q293*)* expression was driven specifically in the epidermis using the *dpy-7* promoter [38]. Here, we observed constitutive expression of the *chil-27p*::GFP reporter indicating that epidermal-specific PALS-25(Q293*) can activate the ORR (Fig 6A). For additional investigation, we developed a construct where *pals-25(Q293*)* expression was driven specifically in the epidermis of adults using a distinct promoter, *col-19p*, which is expressed beginning in early adulthood [39]. The construct was injected into *pals-22 pals-25(jy80)* double mutants, so that the only source of *pals-25* would be from the *col-19p::pals-25(Q293*)* construct (Fig 1A, S2 Table) [19]. Here, in two independently isolated transgenic lines we observed PALS-25(Q293*)-mediated induction of *pals-5p::*GFP expression in young adult animals, but not in L4 stage animals, indicating that the *col-19p::pals-25(Q293*)* construct was functional for inducing the IPR when the promoter is active (Fig 6B). Interestingly, *pals-22 pals-25(jy80); col-19p::pals-25(Q293*)* transgenic animals displayed *pals-5p*::GFP expression not only in the epidermis, but also in the intestine, indicating PALS-25 activation in the epidermis triggered *pals-5p*::GFP expression in the intestine (Fig 6C).

**Fig 6.**
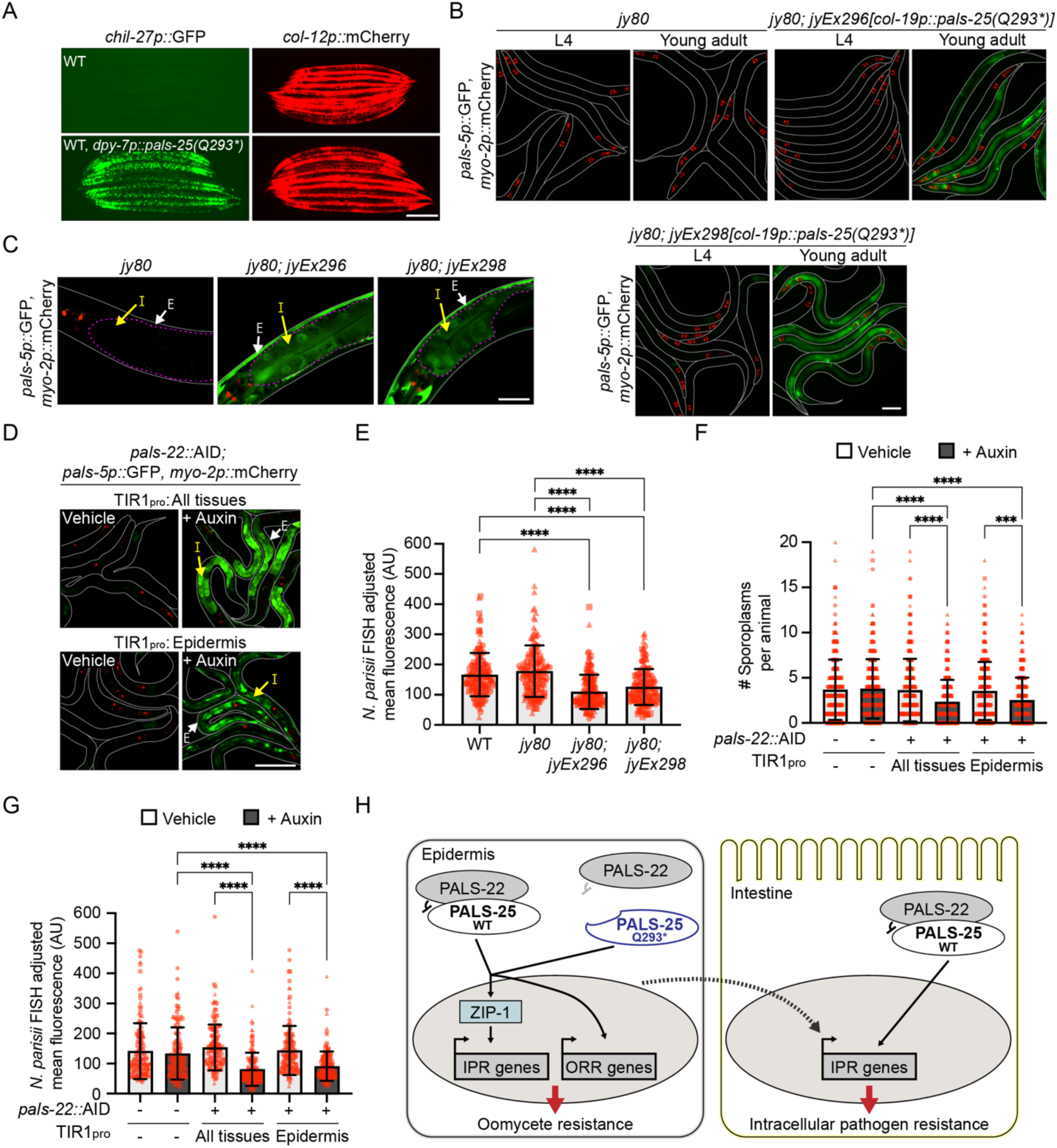
A PALS-22/25-mediated immune response in the epidermis triggers protective immunity in the intestine. **A)** Epidermal (*dpy-7p*)-specific expression of *pals-25(Q293*)* leads to constitutive expression of *chil-27p::*GFP. *col-12p::*mCherry is part of the same transgene and is constitutively expressed starting from the L4 stage. L4 and young adult animals are shown. Scale bar = 100 µm. **B)** Adult-specific expression of *pals-25(Q293*)* in the epidermis induces *pals-5p::*GFP expression. Staged L4 animals and young adult are shown for each genotype. Scale bar = 100 µm. **C)** Higher magnification imaging shows that epidermis-specific expression of *pals-25(Q293*)* in young adults induces *pals-5p::*GFP expression in both the epidermis (white arrows ‘E’) and intestine (yellow arrows ‘I’). Scale bar = 20 µm. Exposure time was set to optimize intestinal *pals-5p::*GFP expression (dashed magenta lines) **D)** Auxin-mediated depletion of PALS-22 in all tissues, or specifically in the epidermis, induces *pals-5p::*GFP expression in both the epidermis (white arrows ‘E’) and intestine (yellow arrows ‘I’). L1 animals treated with either auxin or vehicle control for 24 h at 20 °C are shown. Scale bar = 50 µm. **E)** Epidermis-specific expression of *pals-25(Q293*)* results in increased resistance to *N. parisii* at 30 hpi of young adult animals compared to both WT and a *pals-22 pals-25(jy80)* background without the *pals-25(Q293*)* transgene. Kruskal-Wallis test with Dunn’s multiple comparisons test. n = 150 animals per genotype, three experimental replicates. Symbols represent *N. parisii* FISH fluorescence signal normalized to body area for individual worms; head regions excluded because of expression of the *myo-2p::*mCherry co-injection marker. **F-G)** Auxin-mediated depletion of PALS-22 in all tissues, or specifically in the epidermis, increases resistance to *N. parisii* at 3 hpi **(F)** or 30 hpi **(G)** when compared to vehicle control. Two-way ANOVA with Sidak’s multiple comparisons test. 3 hpi: n = 300 animals per genotype, three experimental replicates. Symbols represent the number of *N. parisii* sporoplasms that infected an individual animal, as determined by *N. parisii*-specific fluorescent FISH signal. 30 hpi: n = 150 animals per genotype, three experimental replicates. Symbols represent *N. parisii* FISH fluorescence signal normalized to body area for individual worms; head regions excluded because of expression of the *myo-2p::*mCherry co-injection marker. For **E-G**, bar heights indicate mean values and error bars represent standard deviations. Different symbol shapes represent animals from infections performed on different days. **** *p* < 0.0001, *** *p* < 0.001. **H)** Model for PALS-22/25-mediated activation of the IPR and ORR. PALS-25(WT) is normally repressed by PALS-22; in *pals-22* mutants the IPR and ORR are induced and mutant animals are resistant to natural pathogens. In *pals-25(Q293*)^icb98^* and *pals-25(Q293*)^jy111^* mutants, PALS-25(Q293*) is constitutively released from repression by PALS-22 to induce the IPR, ORR, and resistance to natural pathogens. PALS-25(WT) or PALS-25(Q293*)-mediated induction of the IPR in the epidermis can signal cell non-autonomously to induce IPR gene expression and intracellular pathogen resistance in the intestine.

Next, as a separate method to activate PALS-25 in an epidermal-specific manner, we performed tissue-specific depletion of its negative regulator PALS-22. Here, we used strains where the endogenous PALS-22 protein was C-terminally tagged with the auxin-inducible degron tag (AID), which causes protein degradation when co-expressed with the TIR1 ubiquitin ligase F-box protein [27, 40]. We used strains where TIR1 was expressed with a ubiquitous promoter (*rpl-28p)* or with an epidermis-specific promoter (*dpy-7p*) [27]. Strains that expressed *rpl-28p::*TIR1*; pals-22::*AID or *dpy-7p::*TIR1*; pals-22::*AID were then crossed into a *pals-5p::*GFP background and exposed to auxin to deplete PALS-22::AID protein, and then analyzed for IPR activation in different tissues. For both ubiquitous and epidermis-specific depletion of PALS-22 we observed *pals-5p*::GFP expression in both epidermal and intestinal tissues of L1 animals after 24 h auxin treatment, and no *pals-5p*::GFP expression on vehicle control (Fig 6D and Materials and Methods). Together, our results from epidermis-specific PALS-25(Q293*) expression and auxin-mediated depletion of PALS-22 indicate that PALS-25 activity in the epidermis can signal to the intestine to induce the *pals-5p::*GFP IPR reporter.

Finally, we assessed if activation of PALS-25 in the epidermis alone could promote resistance to infection in the intestine. For *col-19p::pals-25(Q293*)* transgenic strains where PALS-25(Q293*) was expressed in the epidermis, we infected animals with the intestinal intracellular pathogen *N. parisii* and assessed pathogen load. At 30 hpi, we found that both of the transgenic lines tested were resistant to *N. parisii* when compared to wild-type animals, and when compared to the *pals-22 pals-25(jy80)* double mutants, which have resistance similar to wild-type (Fig 6E and S8A Fig) [19]. For auxin-mediated depletion of PALS-22 in all tissues, or specifically in the epidermis, we found that auxin-treated animals were resistant to *N. parisii* when compared to control animals at both 3 hpi and 30 hpi (Fig 6F, G and S8B Fig). Thus, activation of PALS-25 in the epidermis can induce systemic immune responses to promote resistance against a natural pathogen of the intestine.

## Discussion

In this study, we identify and characterize a gain-of-function allele of *pals-25 (Q293*)*, which activates epidermal and intestinal epithelial defense against natural pathogens of *C. elegans*. Previously isolated loss-of-function mutations of *pals-25* only had an effect in a *pals-22* mutant background, not in a wild-type background [19]. Here, we observe that truncation of the C-terminus of PALS-25 protein by 13 amino acids is sufficient to induce the ORR/IPR defense programs even in the presence of the wild-type negative regulator PALS-22 (Figs 1-3). Loss of the C-terminal 13 amino acids causes PALS-25 protein to lose its normal association with PALS-22, providing a potential explanation for PALS-25(Q293*) phenotypes in a wild-type background (Fig 4). Activation of PALS-25 promotes nuclear localization of ZIP-1 predominantly in the epidermis (Fig 5), yet is able to promote resistance to *N. parisii* and Orsay virus in the intestine, which we found is due to PALS-25 activation in the epidermis signaling cell non-autonomously to upregulate defense in the intestine (Fig 6H). This finding represents a novel example of cross-talk between peripheral tissues providing protective immunity in *C. elegans*.

We find that ORR/IPR gene expression is only partially upregulated in *pals-25(Q293*)* mutants (Fig 2). Partial induction is associated with increased pathogen resistance, but not increased thermotolerance or growth defects observed in *pals-22* mutants (Fig 3). The increased thermotolerance of *pals-22* mutants involves the upregulation of a multi-subunit cullin-RING ubiquitin ligase that includes the cullin CUL-6, RING domain protein RCS-1, functionally redundant Skp-related proteins SKR-3, SKR-4, and SKR-5, and non-redundant F-box proteins FBXA-158 and FBXA-75 [35]. All of the aforementioned components are regulated by wild-type *pals-22/25*. However, in *pals-25(Q293*)* mutants we did not observe upregulation of *fbxa-75*, and there was only modest upregulation of other ubiquitin ligase components, perhaps explaining why these mutants are not thermotolerant (Fig 3E, S2 Fig). Future studies will be aimed at identifying players that interact with PALS-22 and/or PALS-25 proteins and may be responsible for regulating expression of distinct IPR genes and the variety of phenotypes associated with this immune program.

Our co-IP/WB analysis in *C. elegans* together with our analysis in the yeast two-hybrid system reveals that PALS-22 and PALS-25(WT) are physically associated, while the truncated PALS-25(Q293*) protein loses this association with PALS-22 (Fig 4). Together, these results support a model where PALS-22 physically represses the ability of PALS-25 to activate downstream signaling, and the interaction of the two proteins requires the C-terminus of PALS-25. Biochemical functions and protein structures have yet to be determined for any of the *pals* genes, but given that the ORR/IPR is activated upon separation of PALS-22 and PALS-25 proteins, it is intriguing to hypothesize that PALS-22/25 may act as a molecular ‘tripwire’ [41] via an association-dissociation mechanism. In such a scenario the pair would dissociate in response to specific stimuli, such as a pathogen virulence factor, and release PALS-25 to induce the IPR. Here, PALS-22 and PALS-25 share intriguing similarities with certain plant “R gene pairs,” which encode paralog proteins that bind tightly to each other and direct opposite effects on immunity [42, 43]. After sensing virulence factors from a co-evolved plant pathogen, the negative regulator of the R pair will activate immunity by de-repressing the positive regulator of the R pair. In these cases however, the two R gene proteins remain physically associated during the activation phase [44], although dissociation or loss of the negative regulator will activate immunity, like dissociation from or loss of PALS-22 that we find in *C. elegans* [42, 43]. Similarly, the activation of PALS-25 may thus be triggered by currently unknown pathogen-derived factors, which could lead to its dissociation, or if it is similar to R gene proteins, its activation while still binding its negative regulator. Notably however, our previous results show that *pals-22/25* are dispensable for ORR/IPR activation by oomycetes, *N. parisii*, Orsay virus, proteasome inhibition, and other known ORR/IPR activators [18, 19]. Therefore, either PALS-22/25 do not detect these activators, or they detect these activators redundantly with other factors.

As a relative of the ‘tripwire’ model, PALS-22/25 may act together to carry out a shared molecular function that is disrupted during infection. In this scenario, *pals-22* or *pals-25(Q293*)* mutants have impaired cellular function, perhaps due to “free” PALS-25. The ORR/IPR may thus be activated downstream of such a ‘homeostasis altering molecular process’ (HAMP) [45]. There are several examples of IPR induction as a consequence of altered cellular homeostasis. Disruption of protein homeostasis by prolonged heat stress, or by genetic and pharmacological inhibition of proteasome function, activates the IPR [19, 26, 28, 29]. Furthermore, a putative IPR-regulating HAMP was recently identified with mutation of the purine nucleoside phosphorylase *pnp-1,* which causes perturbations in purine metabolism and activation of the IPR [32].

To our surprise, we found that epidermal-specific activation of PALS-25 also activates the IPR in the intestine, indicating that *C. elegans* can coordinate immune responses across the ‘skin-gut’ axis (Fig 6). While there are many examples of signaling from the nervous system to peripheral tissues, less is known about signaling between peripheral tissues like the epidermis and intestine, although findings in a variety of organisms indicate there is such communication. For example, *Drosophila melanogaster* necrotic inflammation of epidermal wing cells activates the innate immune IMD pathway in the intestine [46]. In rats and humans, UV irradiation or burn injury to the skin has been shown to alter gastrointestinal hyperactive immune states [47, 48]. In *C. elegans*, epidermally-expressed *lin-26* regulates expression levels of five distinct fluorescent transcriptional reporters in the intestine [49]. In the context of skin-to-gut immune signaling, Zugasti *et al.* demonstrated that epidermal-specific knockdown of the VH1 dual-specificity phosphatase family gene *vhp-1*, a negative regulator of the p38 MAPK pathway, resulted in intestinal *nlp-29p::*GFP reporter expression [50]. This study also showed that activation of the mitochondria unfolded protein response in the intestine can alter *nlp* gene expression in the epidermis [50]. Signaling across the nematode ‘skin-gut’ axis has also been shown to regulate development and metabolism. Specifically, disruption of epidermal microRNA expression can impact mTORC2 signaling in the intestine to alter vitellogenin (*vit)* gene expression [51]. However, in all of the cases mentioned above, the signals responsible for cross-talk along the ‘skin-gut’ axis are unknown. In a highly mechanistic study of ‘skin-gut’ communication where a secreted signal was identified, Lin and Wang demonstrated that the nuclear hormone receptor NHR-25 expressed in the epidermis regulates the expression and secretion of Hedgehog-like protein GRL-21, which binds to the Patched receptor PTR-24 expressed in the intestine to regulate mitochondria fragmentation and lipid homeostasis in that tissue [52]. In the case of the PALS-25 epidermal-to-intestine IPR immune program described here, we do not know the nature of the signal, nor whether the signaling is direct (i.e. there may be a neuronal intermediate between skin and gut), but our findings add to a growing list of intriguing examples of peripheral tissue communication in *C. elegans*. Future work dissecting the ‘skin-gut’ axis of the IPR may shed light on new mechanisms of inter-tissue coordination of immune responses.

## Materials and Methods

### *C. elegans* maintenance

Worms were maintained using standard methods at 20 °C on Nematode Growth Media (NGM) agar plates top plated with streptomycin-resistant *Escherichia coli* OP50-1, unless stated otherwise [53]. Worm strains used in this study are listed in S1 Table.

### Synchronization of *C. elegans*

To obtain synchronized populations, gravid adult animals were collected from NGM+OP50-1 plates in M9 media into a 15 ml conical tube. Tubes were centrifuged and the supernatant was removed leaving ∼2 ml of M9. 800 µl of 5.65-6% sodium hypochlorite solution and 200 µl of 2 M NaOH were then added to the tube for ∼ 2 min. Embryos released after bleaching were suspended in 15 mL M9, centrifuged, and the supernatant was removed. The wash-and-centrifuge procedure was repeated 5 times and embryos were resuspended in a final volume of 5 ml of M9. Embryos were placed in a 20 °C incubator with continual rotation for 16-24 h to hatch L1s.

### EMS mutagenesis and mapping *pals-25(Q293*)^icb98^*

L4 stage *C. elegans* carrying the *icbIs4[chil-27p*::*gfp*, *col-12p*::*mCherry]* transgene were treated with 24 mM ethyl methanesulfonate (EMS) (Sigma-Aldrich, St. Louis, MO) using previously described procedures [54]. F2 animals constitutively expressing *chil-27p::*GFP were manually picked using a Axio ZoomV16 dissecting scope (Zeiss, Jena, Germany). The GFP positive mutant was crossed with the polymorphic Hawaiian isolate CB4856, and whole genome sequencing data from GFP positive F2 recombinants was analyzed using the CloudMap pipeline [55] to identify the *pals-25(Q293*)^icb98^* allele.

### CRISPR/Cas9-mediated mutagenesis to generate *pals-25(Q293*)^jy111^*

CRISPR/Cas9-mediated mutagenesis with homology directed repair to introduce the Q293* mutation in *pals-25* was performed using a co-CRISPR strategy as previously described [56]. We designed a crRNA overlapping Q293 as well as a repair template to alter the CAA codon to a TAA stop (S2 Table). The crRNA and repair template were synthesized as oligonucleotides (Integrated DNA Technologies, Coralville, IA) and were used with a crRNA targeting the *dpy-10* gene. 67 µM *pals-25* crRNA, 17 µM *dpy-10* crRNA, and 40 µM of commercial tracrRNA were combined and incubated for 5 min at 95 °C followed by 5 min at room temperature. After annealing, the resulting gRNA mixture was added to Cas9-NLS to a final concentration of 27 µM Cas9 (QB3, Berkely, CA) and 8 µM of repair template. *jyIs8[pals-5p::gfp, myo-2p::mCherry]* worms were microinjected and Dpy F1 progeny were screened by PCR for successful introduction of the CAA to TAA early stop mutation. A selected line was crossed out of the Dpy background and backcrossed three times to *jyIs8* animals to generate ERT738 (*pals-25(jy111)*; *jyIs8[pals-5p::gfp, myo-2p::mCherry]*) before experimental testing. ERT738 animals were then backcrossed once to WT to generate ERT751 (*pals-25(jy111)*) (S1 Table).

### Molecular cloning and transgenesis

The *pals-25(Q293*)^icb98^* allele was amplified using the dpy-7p25F/p25ch7R primer pair and inserted via Gibson assembly into pIR6 (*dpy-7p::unc-54 3’utr)* to generate the plasmid pCV6 (*dpy-7p::pals-25(Q293*)::unc-54*). pCV6 was injected into MBA281 *icbIs4[chil-27p*::*gfp*, *col-12p*::*mCherry]* at 50 ng/µl animals along with *myo-2p::dsRed* at 5 ng/µl as the co-injection marker. The *pals-25(Q293*)^jy111^* allele tagged with wrmScarlet modified with synthetic introns was synthesized as a HiFi gBlock (Integrated DNA Technologies) with 50 bp 5’ homology to the *col-19* promoter region and 50 bp homology with the *unc-54* 3’utr. 665 bp of the *col-19p* was PCR-amplified from WT genomic DNA, and 700 bp of the *unc-54* 3’utr was PCR-amplified from pET631 [*pals-22p::pals-22::gfp*::*SBP_3xFLAG::unc-54 3’utr, unc-119(+)*] using GoTaq Green Master Mix (Promega, Madison, WI). The *col-19p*, *Q293*^jy111^* gBlock, and *unc-54* 3’utr pieces were Gibson assembled using the NEBuilder assembly kit and the assembled product was then PCR-amplified with Phusion DNA polymerase (New England Biolabs, Ipswich, MA). The PCR-amplified assembly was A-tailed for 30 min at 70 °C, ligated into pGEM T-Easy vector (Promega), and transformed into DH5*α* cells (New England Biolabs). The resulting vector, pET729 [*col-19p::pals-25(Q293*)::wrmScarlet::unc-54 3’utr*], was verified by restriction digest and sequencing across the insert. pET729 was injected at 60 ng/µl with pBluescript at 40 ng/µl into a *pals-22 pals-25(jy80); jyIs8*[*pals-5p::gfp, myo-2p::mCherry*] background. Transgenic lines were identified by post-L4 induction of *pals-5p::*GFP expression. Three transgenic lines were isolated from different injected P0 animals and two lines were analyzed in this study. A complete list of constructs can be found in S9 Table. Primers used for molecular cloning are listed in S3 Table.

### RNA interference

RNAi was performed by the feeding method using previously described *pals-22*, *pals-25*, and *zip-1* RNAi clones [19, 26, 28]. *For pals-25(Q293*)^icb98^* analysis (Fig 1), 6-8 L4 stage animals were transferred onto RNAi plates (NGM plates supplemented with 5 mM IPTG and 1 mM carbenicillin) seeded with a 300 µl lawn of *E. coli* HT115 carrying the L4440 plasmid (empty vector control) or the *pals-25(RNAi)* clone. The plates were incubated at 20 °C for 72 h and F1 animals at L4 stage were scored for the expression of *chil-27p::*GFP. For *pals-25(Q293*)^jy111^* analysis, synchronized L1 animals were transferred to RNAi plates seeded with a 300 µl lawn of either empty vector control, *pals-25(RNAi)* (Fig 1), or *zip-1(RNAi)* (Fig 5). The plates were incubated at 20 °C for 44 h and L4 stage animals were scored for the expression of *pals-5p::*GFP*, myo-2p::*mCherry. For ZIP-1::GFP analysis, synchronized populations of L1s for *zip-1::*GFP or *pals-25(Q293*)^jy111^ zip-1::*GFP were transferred to standard NGM OP50-1 plates or RNAi plates seeded with a 300 µl lawn of empty vector control, *pals-22(RNAi)*, or *zip-1(RNAi)* (Fig 5). The plates were incubated at 20 °C for 48 h and young adult stage animals were scored for nuclear ZIP-1::GFP expression in the epidermis and/or intestine on a LSM700 confocal microscope with Zen2010 software (Zeiss).

### RNA isolation, qRT-PCR, and sequencing

*pals-25(Q293*)^icb98^* analysis: Synchronized L4 stage WT *and pals-25(Q293*)^icb98^* mutant animals were collected and RNA was extracted by standard procedure using TRIzol (Life Technologies, Carlsbad, CA). RNA samples were sequenced by Genewiz (Azenta Life Sciences, South Plainfield, NJ). Sequencing reads were aligned to WS235 transcriptome from WormBase and transcript abundance was estimated using Kallisto (S4 Table) [57]. This was followed by differential expression analysis using Sleuth and Wald test for two-sample comparisons (S5 Table) [58]. The data was filtered with pval<0.01, qval<0.1, and log2 fold change > 2 to obtain the final list for differentially expressed genes for Venn diagrams, GSEA, and WormCat analysis. Overlap of datasets was visualised using the webtool http://bioinformatics.psb.ugent.be/webtools/Venn/ followed by statistical analysis using a hypergeometric test (http://nemates.org/MA/progs/overlap_stats.html). Heatmaps were produced using Prism 9 (GraphPad, San Diego, CA).

*pals-25(Q293*)^jy111^* analysis: ∼7,000 synchronized L4 stage worms per genotype were grown on 10-cm NGM plates seeded with OP50-1 and were collected (44 h at 20 °C for all strains, except for *pals-22(jy3)* mutants, which are developmentally delayed [26] and were grown for 48 h at 20 °C). RNA was extracted with TRI reagent (Molecular Research Center, Inc., Cincinnati, OH), isolated with BCP reagent (Molecular Research Center), washed with isopropanol and ethanol, and was resuspended in nuclease free (NF) H_2_O. Samples were further purified with a RNeasy Mini kit (Qiagen, Hilden, Germany) and again resuspended in NF-H_2_O for qRT-PCR and RNAseq. qRT-PCR analysis was performed as previously described [26]. cDNA was synthesized from total RNA using the iScript synthesis kit (Bio-Rad, Herculues, CA). qPCR was performed with iQ SYBR Green Supermix (Bio-Rad) with a CFX Connect Real-Time PCR Detection System (Bio-Rad). Expression values for genes of interest were normalized to the expression of a control gene, *snb-1*, which does not show altered expression following IPR or ORR activation [26]. The Pffafl method was used for quantification [59]. Primers used for qRT-PCR analysis are provided in S3 Table. For RNAseq, RNA quality was assessed with a TapeStation system and cDNA libraries were prepared for paired-end sequencing at the Institute for Genomic Medicine at the University of California, San Diego. Sequencing was performed on a NovaSeq6000 sequencer (Illumina Inc., San Diego, CA). Sequencing reads were aligned to WS235 in RStudio using Rsubread and were quantified using Featurecounts (S4 Table). Differential gene expression was performed using limma-voom on the Galaxy platform (usegalaxy.org) (S5 Table) [60]. Lowly expressed genes (CPM < 0.5) were filtered out and an adjusted *p* value of < 0.05 and log2 fold change > 2 was used to define differentially expressed genes used for Venn diagrams, GSEA, and WormCat analysis. Overlap of datasets, and hypergeometric testing, were performed as described above. Heatmaps were produced using Prism 9 (GraphPad).

### Functional expression analysis

Gene Set Enrichment Analysis (GSEA) software v4.2.3 was used for functional analysis of *pals-25(Q293*)^icb98^* and *pals-25(Q293*)^jy111^* using the GSEA Pre-ranked tool [30]. Differentially expressed genes from RNAseq expression data for *pals-25(Q293*)^icb98^* and *pals-25(Q293*)^jy111^* was converted into a GSEA compatible file, and 93 gene sets for comparison were compiled in Excel and converted into a GSEA compatible file. Independent GSEA was performed for *pals-25(Q293*)^icb98^* and *pals-25(Q293*)^jy111^* vs. wild type controls for each experiment. For both analyses, a signal-to-noise metric of 1000 permutations was used. Of the 93 gene sets, any in which fewer than 15 genes were represented in the RNAseq data from this study were excluded from statistical analysis. A heatmap of the GSEA analysis was produced using Prism 9 (GraphPad). GSEA analysis results are provided in S7 Table. Wormcat analysis of *pals-25(Q293*)^icb98^* and *pals-25(Q293*)^jy111^* was performed with WormCat 2.0 (http://www.wormcat.com/index) using whole genome v2 annotations [31]. WormCat analysis results are provided in S8 Table.

### PALS-22 and PALS-25 protein expression and antibody generation

The full length wild-type *pals-22* and *pals-25* were amplified by PCR through the primers listed in S3 Table. The PCR product were cloned into BsaI-HFv2 digested custom vector derived from pET21a. The resulting plasmids (pBEL1693 and pBEL1692), which include PALS-22 with C-terminal TEV-His-tag and PALS-25 with N-terminal His-TEV-tag respectively, were transformed into Rosetta (DE3) cells (Novagen, Sigma) for protein expression. LB with carbenicillin and chloramphenicol selection was inoculated with Rosetta (DE3)/pBEL1693 or pBEL1692 and grown at 37 °C with shaking at 200 rpm. The overnight culture was diluted 1:50 in LB+carbenicillin/chloramphenicol, then expression was induced by adding IPTG to a final concentration of 1 mM at 16 °C, and allowed to shake overnight. Cells were harvested by centrifugation and resuspended in lysis buffer (50 mM Tris pH 8, 300 mM NaCl, 10 mM imidazole, 2mM DTT, 10% Glycerol, 1 mM phenylmethylsulfonyl fluoride (PMSF)). Cells were lysed using the Emulsiflex-C3 cell disruptor (Avestin, Ottawa, Canada) and then centrifuged at 4 °C, 12,000 g to pellet cell debris. The supernatant, containing soluble PALS-22, was passed twice through NiNTA resin #17531802 (Cytiva, Marlborough, MA), which was then washed with wash buffer (50 mM Tris pH 8, 300 mM NaCl, 40 mM imidazole, 1 mM PMSF, 2 mM DTT, 10% glycerol), and the bound proteins were then eluted in elution buffer (50 mM Tris pH8, 300 mM NaCl, 250 mM imidazole, 1 mM PMSF, 2 mM DTT, 10% glycerol). The fractions containing proteins were collected and further purified by HiLoad 16/600 Superdex 200 pg size-exclusion column #28-9893-35 (Cytiva) in PBS (137 mM NaCl, 2.7 mM KCl, 1.5 mM KH_2_PO_4_, 8.1 mM Na_2_HPO_4_, pH 7.5). The pellet, containing a large amount of insoluble PALS-22 or PALS-25, was resuspended in urea lysis buffer (100 mM NaH_2_PO_4_/10 mM Tris base, 10 mM Imidazole, 8 M Urea [titrated to pH 8 by NaOH]). The solubilized pellet was centrifuged at 4000 g, and the supernatant collected. PALS-22 or PALS-25 from the resulting supernatant was passed twice through NiNTA resin (Cytiva #17531802), which was then washed with urea wash buffer (100 mM NaH_2_PO_4_/10 mM Tris base, 40 mM imidazole, 8 M urea [titrated to pH 8 with NaOH]), and the bound proteins were then eluted in urea elution buffer (100 mM NaH_2_PO_4_/10 mM Tris base, 300 mM imidazole, 8 M urea [titrated to pH 8 by NaOH]). Fractions containing PALS-22 or PALS-25 were pooled, concentrated and dialyzed into dialysis buffer (PBS (137 mM NaCl, 2.7 mM KCl, 1.5mM KH_2_PO_4_, 8.1 mM Na_2_HPO_4_), 3.9 M urea [titrated to pH 8 by NaOH]) overnight at room temperature. The following day, the sample was dialyzed once again in fresh dialysis buffer for 3 h at room temperature. The dialyzed sample, and sample from size exclusion purification, were supplemented with 10% glycerol, flash frozen in liquid nitrogen for storage, and submitted to for custom antibody production (ProSci Inc., Poway, CA). Rabbits were initially immunized with 200 µg full-length His::TEV tagged PALS-22 or PALS-25 antigen in Freund’s Complete Adjuvant. Rabbits were then subsequently boosted with four separate immunizations of 100 µg antigen in Freund’s Incomplete Adjuvant over a 16-week period. Approximately 25 ml of serum was collected and PALS-22 or PALS-25 polyclonal antibody was purified with an immuno-affinity chromatography column by cross-linking PALS-22 or PALS-25 to cyanogen bromide (CNBr)-activated Sepharose 4B gel. Antibody was eluted from the affinity column in 100 mM glycine buffer pH 2.5, precipitated with polyethylene glycol (PEG), and concentrated in PBS pH 7.4 + 0.02% sodium azide. Antibody concentration was determined by ELISA and used in Western blot analysis described below.

### Coimmunoprecipitation of PALS-22/25 proteins and Western blot analysis

For each sample ∼100,000 synchronized L1 animals were plated on 10 x 10-cm NGM+OP50-1 plates (10,000 L1s per plate) and grown for 72 h at 20 °C. 20x concentrated OP50-1 was top plated as needed during growth to prevent starvation. Worms were washed off the plates in M9, washed twice with ice-cold PBS + 0.1% Tween-20 (PBS-T), and resuspended in PBS-T in a final volume of ∼500 µl. The worms were then frozen dropwise in LN_2_ and transferred to −80 °C for storage prior to protein extraction. Frozen worm pellets were ground into powder with a mortar and pestle pre-chilled with LN_2_. Powder was resuspended in ice-cold lysis buffer (50 mM HEPES, pH 7.4, 1 mM ethylene glycol-bis(β-aminoethyl ether)-N,N,N′,N′-tetraacetic acid (EGTA), 1 mM MgCl_2_, 100 mM KCl, 1% glycerol, 0.05% Nonidet P-40, 0.5 mM dithiothreitol (DTT), with 1 x cOmplete, EDTA-free protease inhibitor tablet per 10 ml lysis buffer (Roche, Basel, Switzerland) and total protein extract was collected. The extract was centrifuged for 15 min at 21,000 g at 4 °C and supernatants were syringe-filtered through a 0.45 µm membrane (Whatman, Maidstone, United Kingdom). Protein concentration was determined using Pierce 660-nm protein assay (Thermo Fisher Scientific, Waltham, MA). Lysate concentrations were adjusted to 1 µg/µl and 1 mg of each sample was mixed with 25 µl of anti-FLAG M2 Affinity Gel (Sigma-Aldrich). Immunoprecipitation was performed at 4 °C with rotation at 10 rpm for 2 h. Agarose resin beads were pelleted by centrifugation at 8000 g for 30 s and the unbound fraction for each sample was collected and stored at −30 °C. The agarose resin beads were washed twice with 1 ml lysis buffer for 5 minutes, and then twice with 1 ml wash buffer (50 mM HEPES, pH 7.4, 1 mM MgCl_2_, 100 mM KCl) with rotation. Beads were resuspended in 50 µl of wash buffer and stored at −30 °C prior to analysis. 15 µl of 4x Laemmli buffer (Bio-Rad) with 200 mM DTT was added, samples were boiled for 3 min at 100 °C, and were centrifuged for 10 min at 21,000 g at 4 °C. Proteins were separated on a 4-15% sodium dodecyl sulfate–polyacrylamide gel electrophoresis (SDS-PAGE) precast gel (Bio-Rad) at room temperature, and transferred onto polyvinylidene difluoride (PVDF) membrane at 4 °C, 100 V, for 1.5 h. Blocking for non-specific binding was performed in blocking buffer (5% nonfat dry milk dissolved in PBS-T) for 2 h at room temperature. PVDF membranes were incubated with primary antibodies diluted in 5 ml blocking buffer overnight at 4 °C with rocking (anti-PALS-25 1:1,000 and anti-tubulin 1:4,000 monoclonal anti-*α*-tubulin produced in mouse (MilliporeSigma, Burlington, MA)). After primary antibody incubation, the PVDF membranes were washed three times in PBS-T, and then were incubated with secondary antibodies conjugated with horseradish peroxidase in 5 ml blocking buffer for 1.25 h at room temperature with rocking (for anti-PALS-25 primary: goat anti-rabbit 1:10,000, For anti-tubulin primary: goat anti-mouse 1:10,000). the PVDF membranes were washed three times in PBS-T, then were treated with enhanced chemiluminescence (ECL) reagent (Cytiva) for 5 min, and imaged using a Chemidoc XRS+ with Image Lab software (Bio-Rad). Unbound fraction and extract input at 1 µg/µl were processed in parallel with IP samples as described above. Quantification of PALS-25 band intensities for IP inputs were compared to the control sample using Image Lab software (Bio-Rad) and each sample was normalized to its tubulin expression levels. SDS-PAGE and Western blot for immunoprecipitation of PALS-22::GFP::3xFLAG (S6 Fig) was performed with anti-PALS-22 antibody following the same procedure as described above for PALS-25.

### Yeast two-hybrid analysis

Yeast two-hybrid (Y2H) screening was performed using the ULTImate Y2H cell-to-cell mating process (Hybrigenics Service, Evry, France). Either the coding sequence of PALS-25(WT) (amino acids 1-305) or PALS-25(Q293*) (amino acid 1-292) were synthesized and were cloned into pB66 downstream of the Gal4 DNA-binding domain sequence. Each construct was independently used as a bait to screen a *C. elegans* mixed-stage cDNA prey library that was generated by random priming and inserted into the pP6 plasmid encoding the Gal4 activation domain. The library was screened using a haploid mating approach, as previously described, with one yeast sex type carrying the bait construct (mata) and the other (mat*α*) carrying individual prey constructs [61]. For PALS-25(WT), 104 million interactions were analyzed and 352 clones were selected for growth on media lacking tryptophan, leucine, and histidine. For PALS-25(Q293*), 128 million interactions were analyzed and 366 clones were selected for growth on media lacking tryptophan, leucine, and histidine. Prey fragments of growth-positive clones were amplified by PCR and sequenced at their 5’ and 3’ junctions to identify corresponding protein products in the NCBI GenBank database using an automated procedure. The confidence of the bait-prey interaction was assessed by Predicted Biological Score (PBS), as previously described [62]. The bait-prey network analysis of A-C level PBS scores (Fig 4) was generated in Cytoscape v3.9.1 (www.cytoscape.org) and modified in Adobe Illustrator (Adobe Inc., San Jose, CA). Complete Y2H results for PALS-25(WT) and PALS-25(Q293*) are provided in S10 Table.

### Oomycete infections

Oomycete infection assay was performed in triplicate at 25 °C as described previously [17]. Briefly, 3 animals infected with *M. humicola* JUo1, at the stage of pearl-like sporangia present throughout the body of the worm, were used as donors to initiate infection by transferring them onto NGM plates seeded with 50 µl lawn of OP50 along with 30 live L4 stage recipient animals (n = 3 independent plates with 30 animals, per condition). After 48 h of pathogen exposure, animals were moved onto new plates devoid of any pathogen and were transferred over 24 h until end of the assay. Dead animals were identified by the presence of sporangia in their body and were removed from the plate, while missing animals were censored. The assay was continued until all animals in at least one condition were dead. The survival curve was plotted using Prism 9 (GraphPad) and log-rank test was used to assess statistical significance.

### Microsporidia infections of *pals-22/25* mutants in L1 animals

*N. parisii* spores were prepared as previously described [33]. 0.5 million spores were mixed with OP50-1, M9, and 1,200 synchronized L1 animals, and the mix was top-plated onto 6-cm NGM plates; at least two replicates (two plates) per genotype were set up per infection assay. Three independent infection assays were set up for each timepoint described below. Animals were infected at 25 °C for 3 h or 30 h before fixation in either 4% paraformaldehyde or 100% acetone. Fixed worms were incubated at 46 °C overnight with FISH probes conjugated to the red Cal Fluor 610 fluorophore that hybridize to *N. parisii* ribosomal RNA (Biosearch Technologies, Hoddeson, United Kingdom). 3 hpi samples were analyzed for *N. parisii* sporoplasms using a AxioImager M1 compound microscope (Zeiss). 30 hpi samples were analyzed for *N. parisii* meronts using a COPAS Biosort machine (Union Biometrica, Holliston, MA) where FISH signal for each worm was normalized to the length of the worm using time-of-flight measurements.

### Epidermal expression of PALS-25(Q293*) and microsporidia infections in adults

1200 synchronized L1s for each genotype were plated to 2 x 10-cm plates seeded with OP50-1 (600 L1s per plate processed in technical duplicate) and were grown at 23 °C for 50 h to the young adult life stage. Young adults were washed off the 10-cm plates and pelleted in ∼50 µl M9. 2 million *N. parisii* spores, OP50-1, and M9 were mixed with the worm pellet and top plated on 6-cm NGM plates. The plates were transferred to 25 °C and pulse-infected for 3 h. The plates were then removed from the incubator, infected worms were washed off the plates, transferred to fresh 6-cm NGM plates pre-seeded with OP50-1, and returned to 25 °C for an additional 27 h. Animals were fixed and incubated with *N. parisii* FISH probes as described above. Samples were analyzed for meronts on an ImageXpress Nano plate reader using the 4x objective (Molecular Devices, LLC, San Jose, CA). The worm area was traced using FIJI software (excluding the head regions due to *myo-2p::*mCherry co-injection marker expression) and the average fluorescence intensity of each worm was quantified with the background fluorescence of the well subtracted. 50 animals per genotype were quantified for each experimental replicate, and three independent infection experiments were performed.

### Auxin-mediated depletion of PALS-22 and microsporidia infections in larvae

Strains for auxin-mediated depletion of PALS-22 either in all tissues or specifically in the epidermis were crossed into a *jyIs8[pals-5p::gfp, myo-2p::mCherry]* background (S1 Table) [27]. Gravid adults were bleached to release embryos, washed five times with M9, and 1,200 embryos per strain were plated on 6-cm NGM plates supplemented with 200 µM auxin or 0.15% ethanol control plates without OP50-1 prepared as previously described [27]. The plates were stored for 24 h at 20 °C to allow embryos to hatch and *pals-5p::*GFP expression in the intestinal and epidermal tissues of starved L1s was assessed on a LSM700 confocal microscope with Zen2010 software (Zeiss). 2 million *N. parisii* spores, OP50-1, and M9 were mixed and top-plated onto the starved L1s treated with auxin or ethanol control. Animals were infected for 3 h or 30 h, fixed, and incubated with *N. parisii* FISH probes as described above. 3 hpi samples were analyzed for sporoplasms as described above. 30 hpi samples were analyzed for meronts on an ImageXpress Nano plate reader using the 4x objective (Molecular Devices, LLC). Worm area was traced using FIJI software (excluding the head regions due to *myo-2p::*mCherr*y* co-injection marker expression) and the average fluorescence intensity of each worm was quantified with the background fluorescence of the well subtracted. 50 animals per genotype or treatment were quantified for each experimental replicate, and three independent infection experiments were performed.

### Orsay virus infections of *pals-22/25* mutants in L1 animals

Orsay virus filtrates were prepared as previously described [29]. A 1:5 dilution of Orsay virus filtrate was mixed with OP50-1, M9, and 1,000 synchronized L1 animals, and the mix was top-plated onto 6-cm NGM plates; at least two technical replicates (two plates) per genotype were set up per infection assay. Four independent infection assays were performed. Animals were infected at 20 °C for 18 h before fixation in 4% paraformaldehyde. Fixed worms were incubated at 46 °C overnight with two FISH probes, mixed equally, that were both conjugated to the red Cal Fluor 610 fluorophore and hybridize to either Orsay RNA1 or RNA2 genome segments (Biosearch Technologies). Samples were analyzed for Orsay virus infection using a AxioImager M1 compound microscope (Zeiss). 100 animals per genotype, per infection assay, were scored and the percentage of infected animals in the population was quantified.

### Bead feeding assays

1,200 synchronized L1 animals were mixed with OP50-1, M9, and 6 µl of Fluoresbrite Polychromatic Red Microspheres (Polysciences Inc., Warrington, PA) and the mix was top-plated onto 6-cm NGM plates. Animals were allowed to feed on the fluorescent beads for 5 min at room temperature while the NGM plates dried, followed by 5 minutes at 25 °C. The plates were then immediately moved to ice to reduce feeding rate, washed twice with ice-cold PBS-T, and then were fixed in 4% paraformaldehyde. Fixed worms were then imaged in a 96-well plate on an ImageXpress Nano plate reader using the 4x objective (Molecular Devices, LLC). The worm area was traced using FIJI software and the average fluorescence intensity of each worm was quantified with the background fluorescence of the well subtracted. 50 animals per genotype were quantified for each experimental replicate and three independent experimental replicates were performed. *eat-2(ad465)* mutants were included in the assay as a feeding-defective control [63].

### Thermotolerance assays

Thermotolerance assays were performed as previously described [35]. Briefly, mixed-stage well-fed animals for each genotype were grown at 20 °C. L4 animals were picked off mixed-stage plates and were transferred onto fresh NGM plates seeded with OP50-1; 30 L4s per plate, three plates per genotype, three independent experimental replicates performed on different days. The plates were then subject to heat shock on a single layer for 2 h at 37.5 °C in a dry incubator. The plates were removed from the incubator, placed on a single layer for 30 min at room temperature, and then transferred to 20 °C. After 24 h, survival of each worm was scored as alive or dead, with dead worms assessed as failure to respond to anterior or posterior touch with a worm pick. The percentage of worms alive for each plate was quantified for each genotype. Missing animals were censored from the analysis.

### Development rate measurements

40 gravid adults for each genotype (60 adults for *pals-22(jy3)* due to smaller brood size in this mutant [26]) were transferred onto a 10-cm NGM plate seeded with OP50-1 and were incubated at 20 °C for 2 h. The adults were gently suspended in M9 media and suctioned off the plates while eggs remained embedded in the OP50-1 lawn. The plates were then incubated at 20 °C, and the percentage of animals that had reached L4 stage or older at 48 h or 64 h was quantified. 100 animals per genotype were scored at each timepoint, and three experimental replicates were performed.

### Bortezomib treatment

Inhibition of the proteasome in *zip-1::gfp* animals was performed with bortezomib (Selleckchem Chemicals LLC, Houston, TX) as previously described [28]. Synchronized L1 animals were plated on 6-cm NGM plates seeded with OP50-1 and were grown for 44 h at 20 °C. 10 mM bortezomib in DMSO was top-plated to a final concentration of 20 µM. As controls, the same volume of DMSO vehicle was added to control plates, or plates were left untreated. The plates were incubated for 4 h at 20 °C. Nuclear ZIP-1::GFP expression in the epidermis and/or intestine was analyzed an imaged on a LSM700 confocal microscope with Zen2010 software (Zeiss).

## Acknowledgments

We thank Vladimir Lažetić, Lakshmi Batachari, Cheng-Ju Kuo, and Michael Blanchard for helpful comments on the manuscript. We thank Gira Bhabha for her help with PALS-22 and PALS-25 protein synthesis and for helpful advice on the project. RNAseq data were generated at the UC San Diego IGM Genomics Center utilizing an Illumina NovaSeq 6000 that was purchased with funding from the National Institutes of Health SIG grant (#S10 OD026929). Some strains used in this study were provided by the Caenorhabditis Genetics Center (CGC), which is funded by NIH Office of Research Infrastructure Programs (P40 OD010440).

## Supporting Information Figures and Legends

**S1 Fig.**
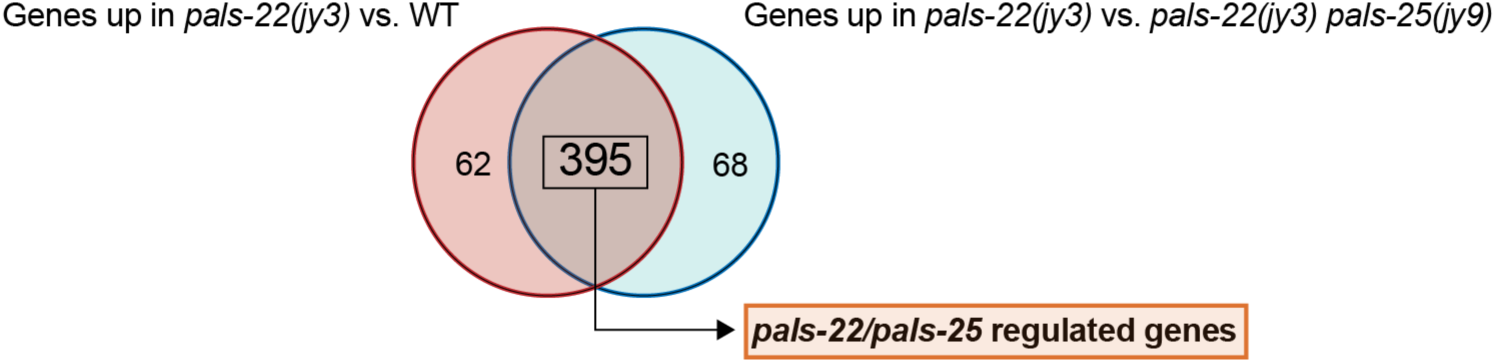
Genes regulated by *pals-22* and *pals-25*. Venn diagram of differentially expressed gene in *pals-22(jy3)* mutants as compared to wild-type animals, and *pals-22(jy3)* mutants as compared to *pals-22(jy3) pals-25(jy9)* double mutants. The overlap of these differentially expressed gene sets is significant and represents genes regulated by *pals-22* and *pals-25*. Hypergeometric test, RF = 26.9; *p* < 0.000e-00.

**S2 Fig.**
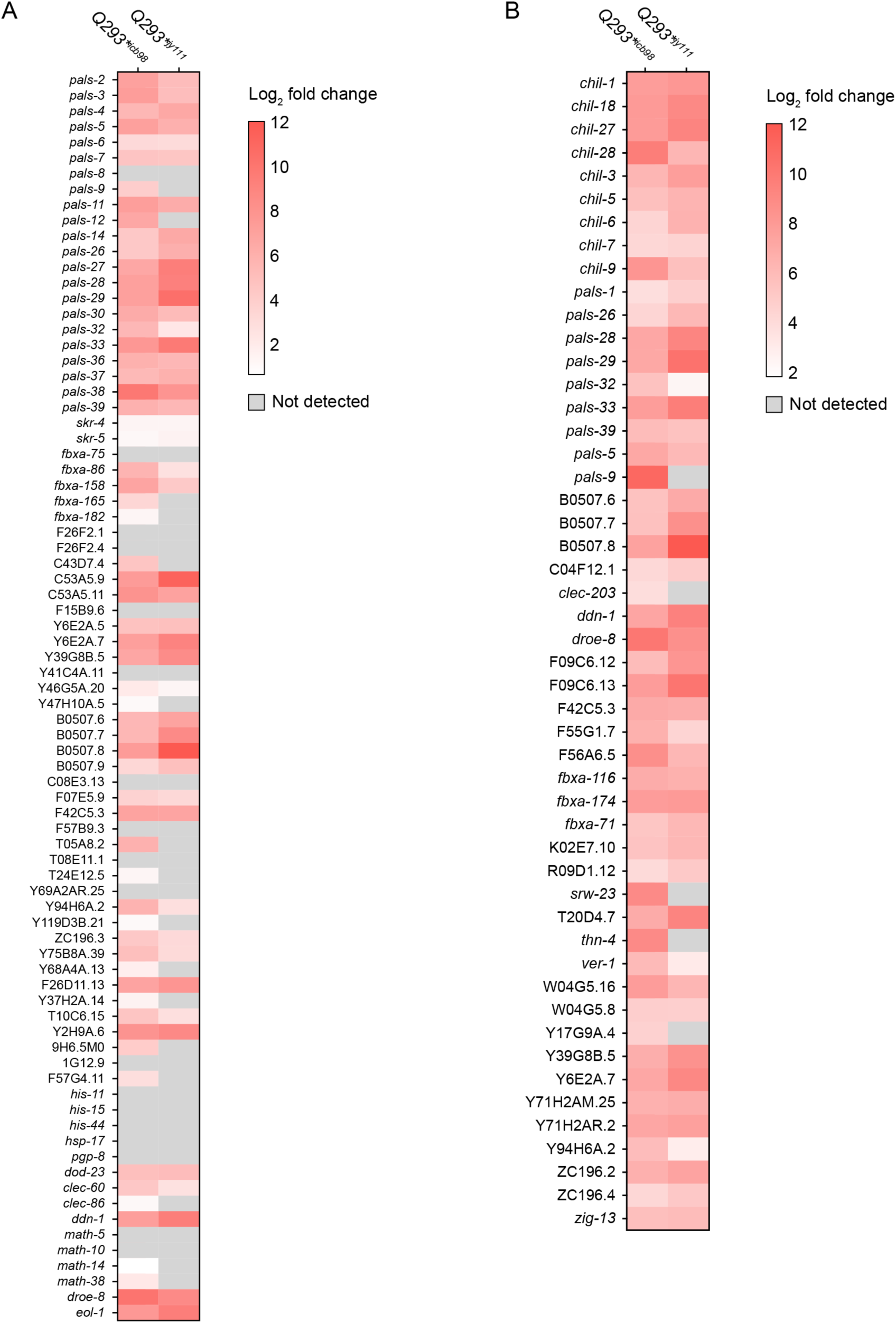
*pals-25(Q293*)* mutants have upregulated genes in common with transcriptional responses to natural pathogens. Heatmap of *pals-25(Q293*)^icb98^* and *pals-25(Q293*)^jy111^* mutant gene expression values measured as log_2_ fold change compared to the 80 genes of the IPR **(A)** and 50 commonly upregulated genes of the ORR **(B)**. Several genes, for example *pals-26*, belong to both the IPR and ORR. For comparisons to the IPR in **A**, some log_2_ fold change values shown for *pals-25(Q293*)^icb98^* and *pals-25(Q293*)^jy111^* mutants are less than 2 for consistency with the previously published Reddy *et al.* 2019 IPR dataset. Genes where transcripts were not detected, or abundances were below cutoff levels during analysis (Materials and Methods), are shown in grey.

**S3 Fig.**
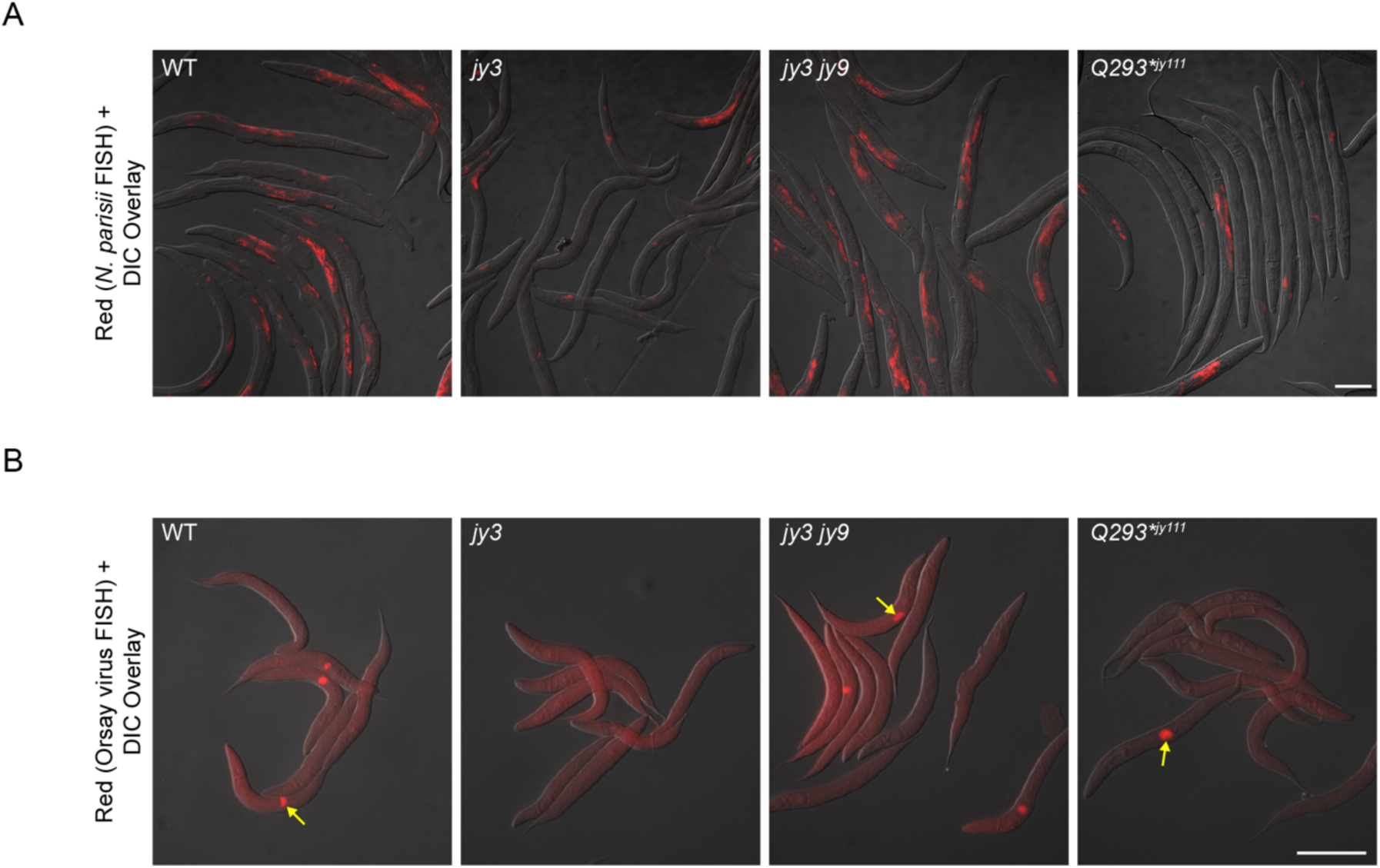
*pals-25(Q293*)^jy111^* mutants display increased resistance to *N. parisii* and Orsay virus. **A)** Representative images of WT animals, *pals-22(jy3)*, *pals-22(jy3) pals-25(jy9)*, and *pals-25(Q293*)^jy111^* mutants infected at L1 with *N. parisii*, fixed at 30 hpi, and stained with *N. parisii-* specific FISH probe (red fluorescence). **B)** Representative images of WT animals, *pals-22(jy3)*, *pals-22(jy3) pals-25(jy9)*, and *pals-25(Q293*)^jy111^* mutants infected at L1 with Orsay virus, fixed at 18 hpi, and stained with Orsay virus-specific FISH probe (red fluorescence representing viral infection denoted by yellow arrows). For **A, B** scale bar = 100 µm. DIC = differential interference contrast.

**S4 Fig.**
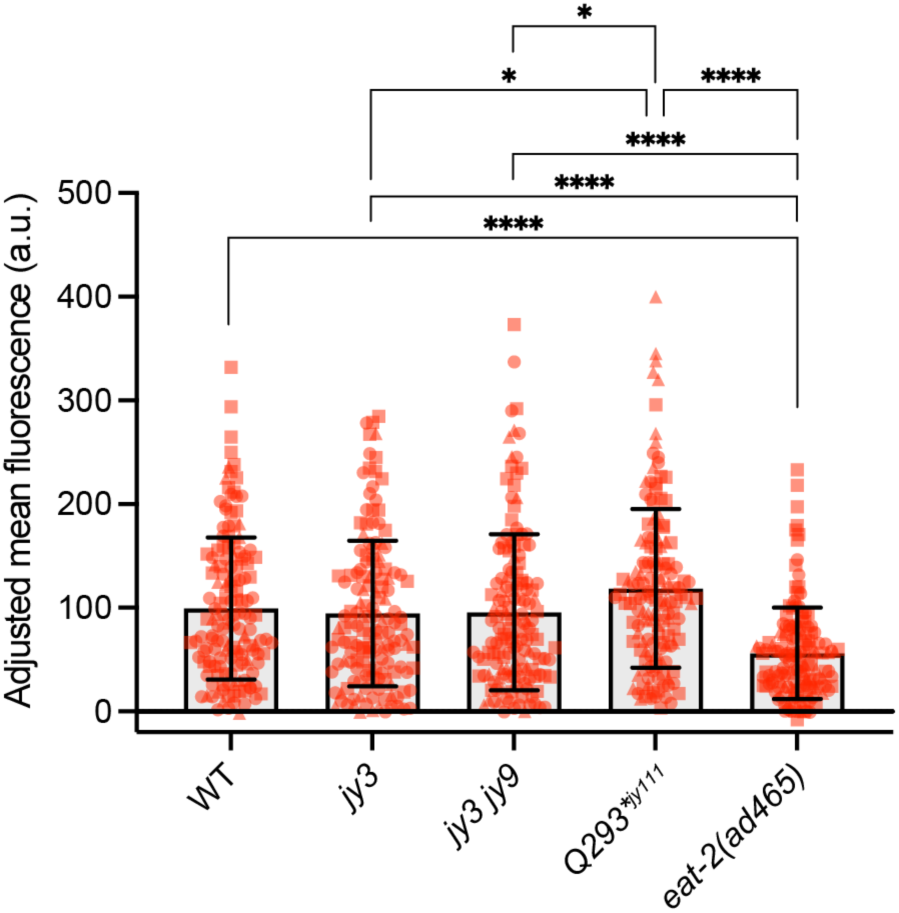
*pals-25(Q293*)^jy111^* mutants do not have lowered accumulation of fluorescent beads in the intestinal lumen. Quantification of fluorescent bead accumulation in WT animals, *pals-22(jy3)*, *pals-22(jy3) pals-25(jy9)*, and *pals-25(Q293*)^jy111^* mutants. *eat-2(ad456)* mutants have a known feeding defect due to abnormal pharyngal pumping and were used as a positive control. **** *p* < 0.0001, * *p* < 0.05, Kruskal-Wallis test with Dunn’s multiple comparisons test. n = 150 animals per genotype, three experimental replicates. Symbols represent fluorescence measurements for individual animals and different symbol shapes represent animals from experimental replicates performed on different days. Bar heights indicate mean values and error bars represent standard deviations.

**S5 Fig.**
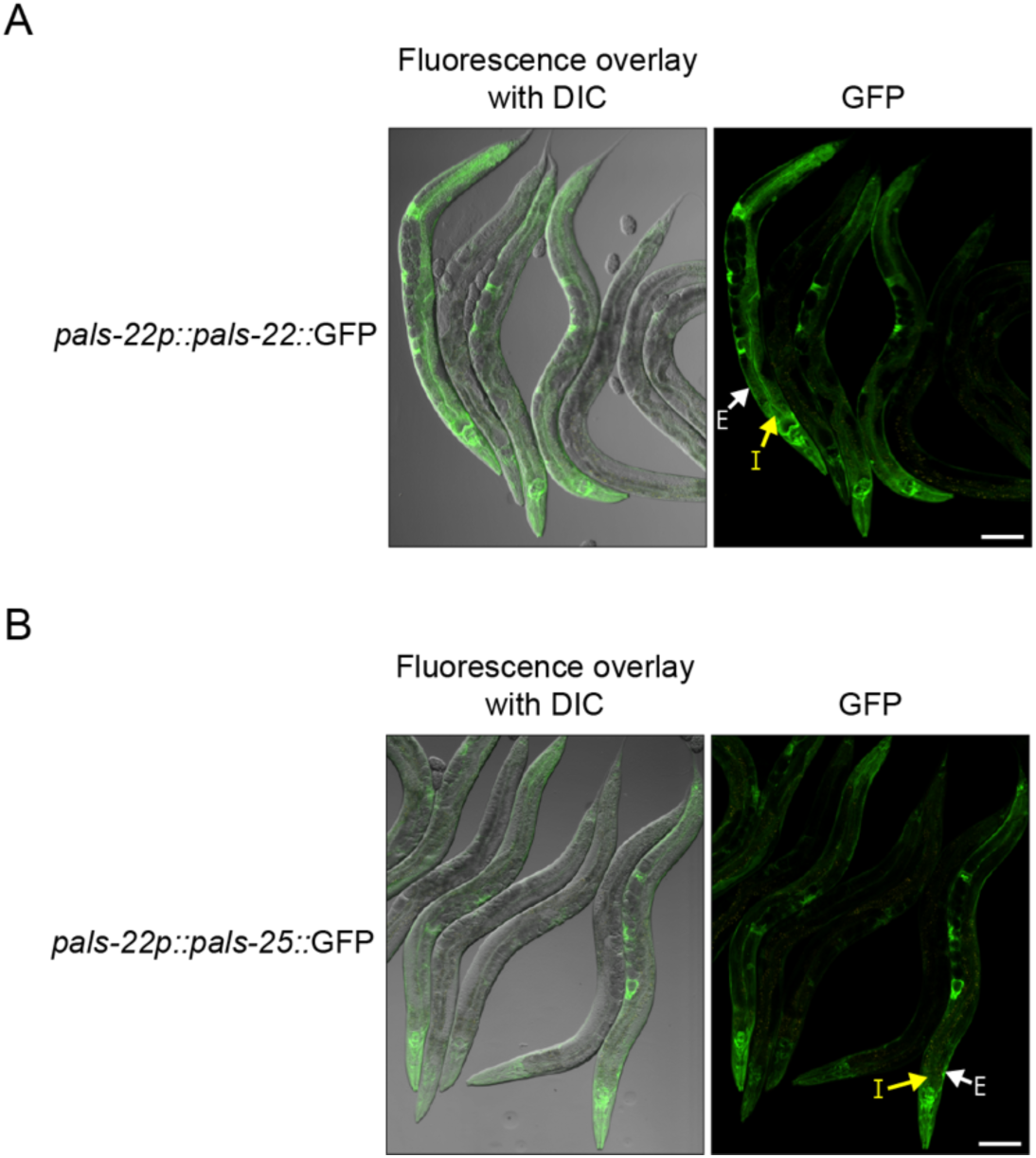
PALS-22::GFP and PALS-25::GFP are broadly expressed. Confocal fluorescence images of adult animals carrying a multicopy TransgeneOme fosmid expressing PALS-22::GFP **(A)** or PALS-25::GFP **(B)**. Fosmids have each gene tagged at the C terminus with GFP, surrounded by approximately 20 kb of endogenous regulatory region. Here, PALS-22 and PALS-25 are expressed in intestinal, epidermal, pharyngal and neuronal tissues. Yellow arrows indicate intestinal tissue (I) and white arrows indicate epidermal tissue (E). Scale bar = 100 µm. DIC = differential interference contrast.

**S6 Fig.**
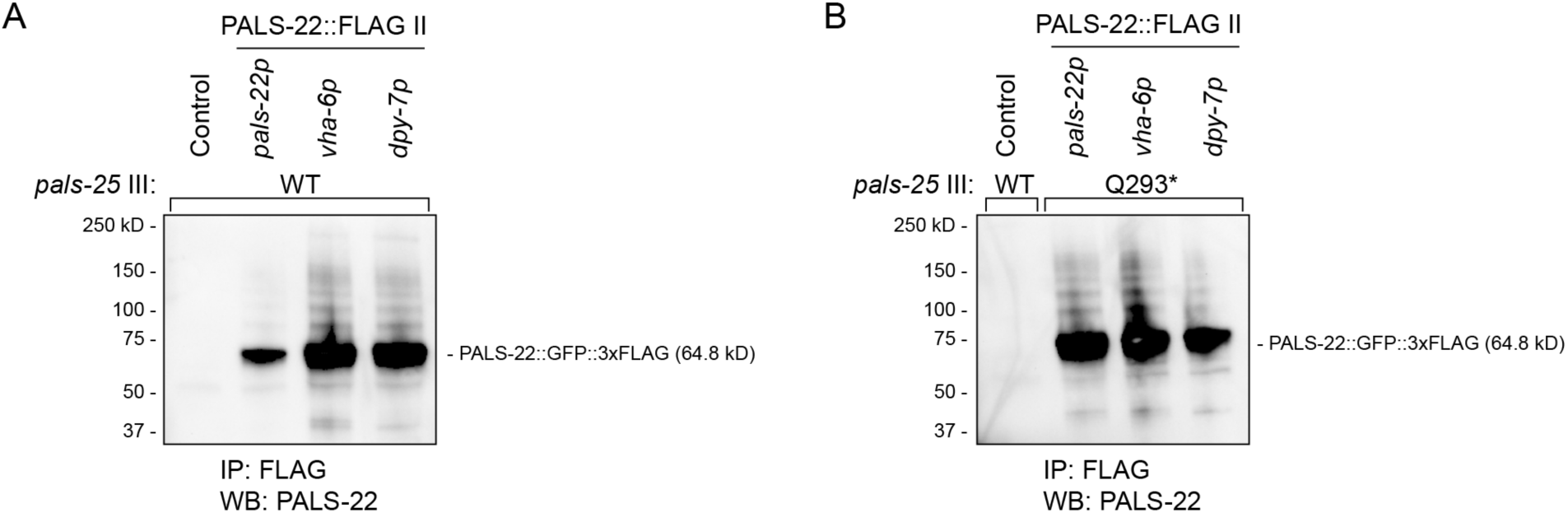
PALS-22::FLAG is captured by FLAG immunoprecipitation. Western blot analysis of PALS-22::GFP::3xFLAG expressed from the MosSCI locus on chromosome II using the endogenous promoter (*pals-22p*), an intestinal promoter (*vha-6p*), or epidermal promoter (*dpy-7p*). FLAG-IP successfully captures PALS-22 in both a WT *pals-25* background **(A)** and in a *pals-25(Q293*)^jy111^* mutant background **(B)**. The GFP::3xFLAG control expressed from the MosSCI locus on chromosome II using the intestinal *spp-5p* promoter does not blot for PALS-22. Western blot was performed using anti-PALS-22 antibody.

**S7 Fig.**
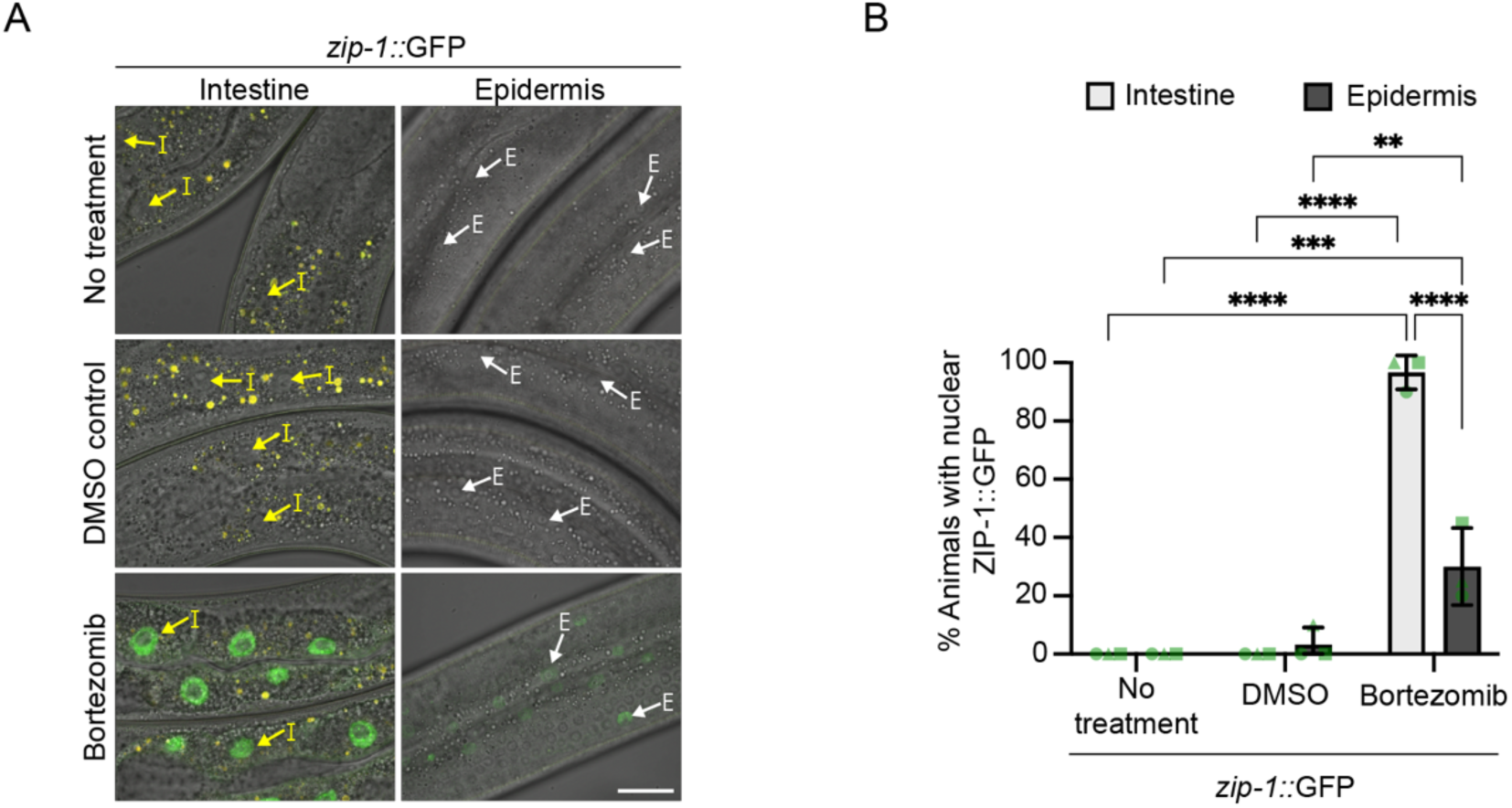
Proteasome inhibition induces nuclear localization of ZIP-1 in the intestine and epidermis. **A)** ZIP-1::GFP is expressed in intestinal and epidermal nuclei following bortezomib treatment, but not in untreated or DMSO control treated worms. Yellow arrows highlight intestinal nuclei ‘I’ and white arrows highlight epidermal nuclei ‘E’. Images are a composite of differential interference contrast, GFP, and RFP fluorescence channels and auto-fluorescent intestinal gut granules appear as yellow signal. Scale bar = 20 µm. **B)** Quantification of **A**, **** *p* < 0.0001, *** *p* < 0.001, ** *p* < 0.01, two-way ANOVA with Sidak’s multiple comparisons test. n = 3 experimental replicates, 20 animals per treatment assessed for both intestinal and epidermal ZIP-1::GFP expression for each replicate. Different symbols represent replicates performed on different days. Bar heights indicate mean values and error bars represent standard deviations.

**S8 Fig.**
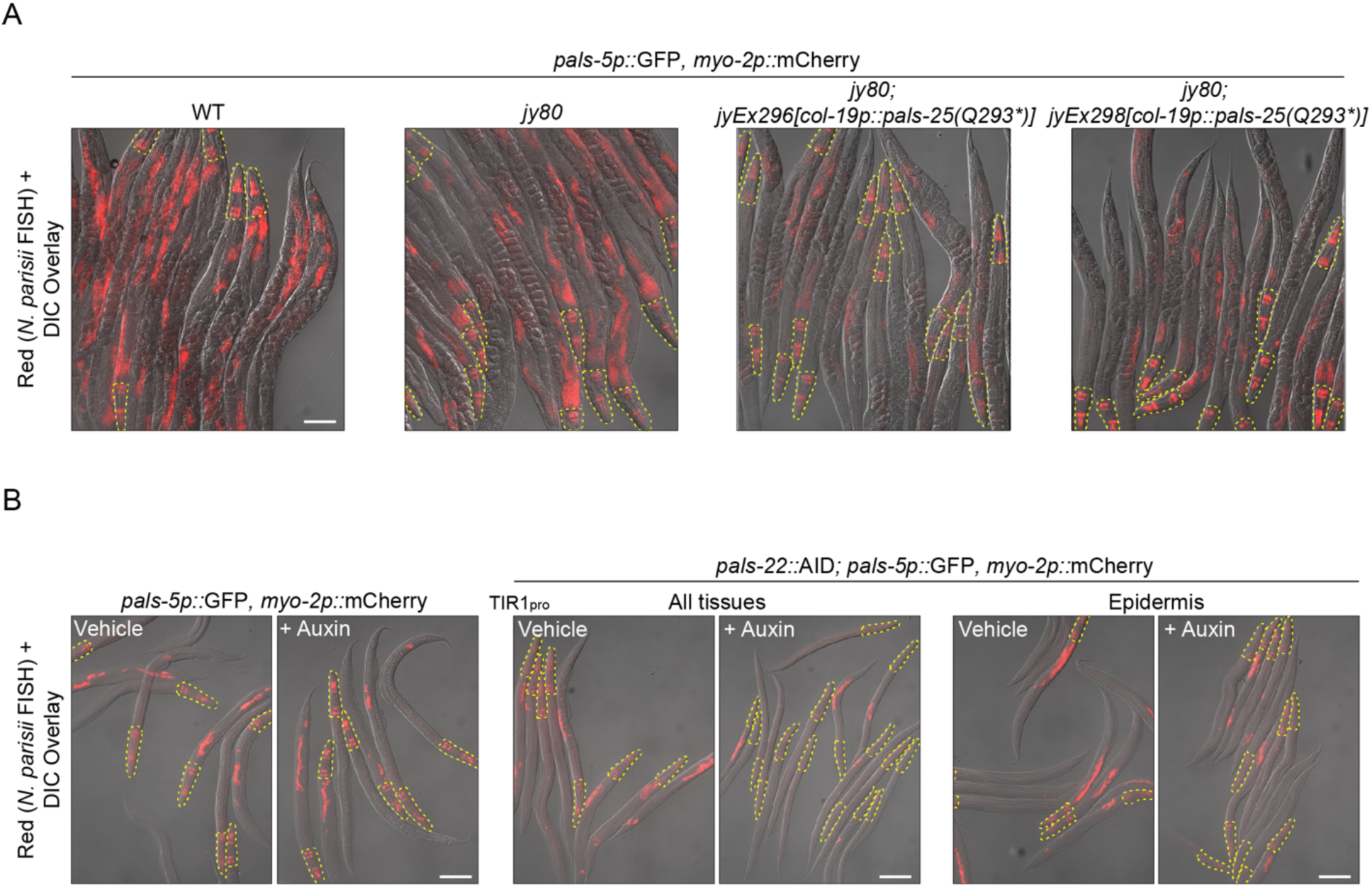
Epidermis-specific expression of *pals-25(Q293\**), or auxin-mediated depletion of PALS-22 in the epidermis, increases resistance to *N. parisii*. **A)** Representative images of WT, *pals-22 pals-25(jy80)*, *pals-22 pals-25(jy80); jyEx296[col-19p::pals-25(Q293*)]*, and *pals-22 pals-25(jy80); jyEx298[col-19p::pals-25(Q293*)]* animals infected as young adults with *N. parisii*, fixed at 30 hpi, and stained with *N. parisii-*specific FISH probe (red fluorescence). **B)** Representative images of WT animals, transgenic animals where PALS-22 can be ubiquitously depleted, or transgenic animals where PALS-22 is epidermally depleted, treated with either vehicle control or auxin. Control or auxin-treated strains were infected at L1 with *N. parisii*, fixed at 30 hpi, and stained with *N. parisii-*specific FISH probe (red fluorescence). For **A, B** scale bar = 100 µm. DIC = differential interference contrast. All strains are in a *jyIs8[pals-5p::gfp, myo-2p::mCherry]* background so head regions (yellow dashed lines) were omitted from pathogen load analysis due to expression of the *myo-2p::*mCherry co-injection marker.

**S1 Table.** Worm strains used in this study.

**S2 Table.** *pals-22/25* mutant alleles and CRISPR reagents used in this study.

**S3 Table.** qPCR primers and cloning primers used in this study.

**S4 Table.** RNAseq statistics. Number of filtered genes = removal of low counts and genes that were undetected by RNAseq. Sanitized = removal of dead genes using the WormBase ‘Gene Name Sanitizer’ function. Significantly upregulated = log2 fold change > 2. Unique = collapse of distinct transcript forms to a single gene.

**S5 Table.** Differentially expressed genes in *pals-25(Q293*)^jy111^*, *pals-25(Q293*)^icb98^*, and *pals-22(jy3)* mutants compared to wild-type animals, and differentially expressed genes in *pals-22(jy3)* mutants compared to *pals-22(jy3) pals-25(jy9)* double mutants.

**S6 Table.** Gene set overlap comparisons and statistics for hypergeometric testing.

**S7 Table.** Gene sets and GSEA results for *pals-25(Q293*)^jy111^* and *pals-25(Q293*)^icb98^*.

**S8 Table.** WormCat analysis of *pals-25(Q293*)^jy111^* and *pals-25(Q293*)^icb98^*. RGS = number of differentially expressed genes from this study in the category. AC = total number of genes in category.

**S9 Table.** Constructs used in this study.

**S10 Table.** Yeast two-hybrid analysis of PALS-25(WT) and PALS-25(Q293*). PBS scores (described in Materials and Methods): A = very high confidence in the interaction, B = high confidence in the interaction, C = good confidence in the interaction, D = moderate confidence in the interaction; can include false-positive interactions and those hardly detectable (i.e. low representation in the mRNA library, improper prey folding, or prey toxicity in yeast), E = interactions involving highly connected (or relatively highly connected) prey domains, may include non-specific interactions, F = experimentally proven technical artifacts (not included).

